# Nhlh1 and Nhlh2, a global transcriptional mechanism regulating commissural axon projection via Robo3 activation

**DOI:** 10.1101/2022.09.23.509112

**Authors:** Aki Masuda, Rieko Ajima, Yumiko Saga, Tatsumi Hirata, Yan Zhu

## Abstract

Commissural neurons are highly heterogeneous in their developmental origins, neurotransmitter type and function, but all share the common feature of projecting axons across the midline. The floor plate-crossing commissural axons in mammals, from the spinal cord to the midbrain, are guided by a conserved molecular mechanism relying primarily on Netrin-1/DCC/Robo3 signaling. Up to date, we know very little about the upstream transcriptional program that specify commissural axon laterality, neither do we know if a common mechanism operates in all commissural neurons. Here, we identified a pair of highly related helix-loop-helix transcription factors, Nhlh1 and Nhlh2, as a global transcriptional mechanism that controls the laterality of all floor plate-crossing commissural axons. Forced expression of Nhlh1/2 induce ectopic Robo3 expression and contralateral axon projections. And mutant mice deficient in both genes show a remarkable reduction in Robo3 expression and a total lack of ventral commissures from the spinal cord to the midbrain. This global mechanism may interact with neuron type specific mechanism to achieve specific generation of commissural circuits.

## INTRODUCTION

In bilaterians, neuronal information must be communicated between the two halves of the central nervous system (CNS) for the normal functions of animals, such as respiration, locomotion, and visual and auditory processing (Borst and Soria van Hoeve, 2012; Bouvier et al., 2010; Goulding, 2009; Kiehn, 2016; Lanuza et al., 2004; Petros et al., 2008). This crucial task is fulfilled by commissural neurons that project their axons across the midline of the neural tube to connect with their target neurons on the contralateral side. The evolution of commissural circuit organization may underlie the emergence of new functions, such as quadrupedal locomotion and binocular vision (Friocourt et al., 2019; Kiehn, 2016; Petros et al., 2008).

Many diverse neuron classes, defined developmentally by distinct combinations of transcription factors (TFs), contain commissural neurons, and each neuron class often contains a mixture of commissural and ipsilaterally-projecting neurons (Alaynick et al., 2011; Lai et al., 2016; Tulloch et al., 2019). Therefore, commissural neurons are highly heterogeneous in terms of their developmental origin, morphology, neurotransmitter type, connectivity, and neurophysiology (Chedotal, 2014; Tulloch et al., 2019), but all share the core defining feature of projecting their axons across the midline. Despite their heterogeneity, axon guidance mechanisms that direct commissural axons to reach and cross the midline, are fairly conserved and well understood (Chedotal, 2011, 2019; Comer et al., 2019). In contrast, the transcriptional programs that specify the laterality of commissural axons, and whether a global regulatory mechanism operates for diverse commissural neurons, are poorly understood.

In vertebrates, most commissural neurons in the spinal cord, hindbrain, and midbrain project ventrally to cross the midline at the floor plate (FP) (Friocourt et al., 2019). The FP does not extend beyond the midbrain, therefore, it is missing from the forebrain (Placzek and Briscoe, 2005). Guidance of commissural axons towards the FP predominantly relies on signaling between the ligand Netrin-1, which is expressed from the FP and the ventral neural progenitors, and its receptor DCC and a facilitator Robo3, which are expressed in the commissural neurons (Dominici et al., 2017; Fazeli et al., 1997; Kennedy et al., 1994; Marillat et al., 2004; Moreno-Bravo et al., 2019; Sabatier et al., 2004; Serafini et al., 1996; Tamada et al., 2008; Varadarajan et al., 2017; Yamauchi et al., 2017). Another ligand receptor pair, Shh and Boc, makes an additional but minor contribution to this process (Charron et al., 2003; Okada et al., 2006; Wu et al., 2019).

Knockout (KO) mice of *Netrin-1*, or *DCC* or *Robo3* showed disrupted ventral commissure formation, among which *Robo3* KO mice showed a complete lack of ventral commissure in the spinal cord and hindbrain (Fazeli et al., 1997; Laumonnerie et al., 2015; Marillat et al., 2004; Sabatier et al., 2004; Serafini et al., 1996; Tamada et al., 2008).

Robo3 was initially proposed to silence the Slit-Robo1/2 mediated repulsion from the midline in pre-crossing commissural axons (Chen et al., 2008; Sabatier et al., 2004). A later study showed that mammalian Robo3 can interact with DCC and facilitate Netrin-1 signaling, hence, directly promoting extension of commissural axons towards the midline (Zelina et al., 2014). Robo3 is transiently expressed in pre-crossing but is quickly downregulated in post-crossing commissural axons/neurons (Marillat et al., 2004; Sabatier et al., 2004), and Robo3 is the only currently known molecule that is exclusively expressed in FP-crossing commissural neurons from the spinal cord to the midbrain (Friocourt et al., 2019; Inamata and Shirasaki, 2014; Marillat et al., 2004; Sabatier et al., 2004; Tamada et al., 2008; Tulloch et al., 2019). DCC, however, shows more broad expression, including some ipsilaterally-projecting neurons (Bai et al., 2011; Dillon et al., 2005; Keino-Masu et al., 1996). Therefore, how Robo3 expression is transcriptionally regulated is likely to be the core of the transcriptional program that specifies commissural identity.

There is limited knowledge on how Robo3 transcription is regulated in commissural neurons. LIM-homeodomain TFs, Lhx2/9, have been shown to control Robo3 expression in the commissural neurons of dI1 spinal neurons, and Lhx2 can bind to the Robo3 promoter (Wilson et al., 2008). However, there is no evidence suggesting that Lhx2/9 can account for Robo3 expression beyond the dI1 spinal neurons. In the midbrain and preBötzinger complex in the hindbrain, neural progenitors expressing the homeobox gene Dbx1 give rise to commissural neurons (Bouvier et al., 2010; Inamata and Shirasaki, 2014). Knockout or knockdown of *Dbx1* affected contralateral axon projections, whereas forced expression of Dbx1 in the midbrain induced ectopic Robo3 expression and contralateral axon projection. However, Dbx1 is also an unlikely solution to account for Robo3 expression beyond those commissural neurons derived from

Dbx1-positive progenitor domains. Moreover, Dbx1, whose expression is confined to neural progenitors, cannot directly regulate the expression of Robo3, which is expressed in young post-mitotic neurons. Therefore, whether Robo3 is separately regulated by different sets of TFs in distinct classes of commissural neurons, or whether there exists a common transcriptional mechanism for all FP-crossing commissural neurons remain unknown. The fact that a highly conserved spatial and temporal distribution of Robo3 has been observed across divergent amniote species indicates that a conserved, possibly common, regulatory program is likely to operate (Friocourt et al., 2019).

In this study, we attempted to uncover transcriptional mechanisms that specify axon laterality in commissural neurons. We reasoned that components of an immediate upstream regulatory program are likely to be enriched in the pre- versus post-crossing commissural neurons, reflecting the temporal dynamics of Robo3 (Marillat et al., 2004; Sabatier et al., 2004). Based on this assumption, we took advantage of the results from our RNA-seq experiment which compared transcriptomes between pre- and post-crossing pontine nucleus (PN) neurons in murine hindbrains, and identified an enrichment of a pair of closely-related basic helix-loop-helix (bHLH) TFs, Nhlh1 and Nhlh2, in the pre-crossing population. Nhlh1 and Nhlh2 (previously known: Nscl1/2, and Hen1/2) are expressed in immature neurons (Duncan et al., 1997; Haire and Chiaramello, 1996; Murdoch et al., 1999; Theodorakis et al., 2002), but their roles in neural development have not been thoroughly explored, particularly in the context of a double deficiency of both genes (Kruger and Braun, 2002; Kruger et al., 2004; Schmid et al., 2007). We found that the *Robo3* promoter contains multiple copies of Nhlh1 and Nhlh2 binding sites, and that forced expression of Nhlh1 and Nhlh2 could induce ectopic Robo3 expression. By generating loss-of-function Nhlh1 and Nhlh2 mouse lines, we were able to show that deficiency in both Nhlh1 and Nhlh2 leads to a marked reduction in Robo3 transcription, and a total disruption of FP-crossing commissures, from the spinal cord to the midbrain.

## RESULTS

To separate relatively pure pre- and post-crossing commissural neurons is not trivial, as during development, the pre- and the post-crossing, as well as the commissural and ipsilateral-projecting neurons, intermingle extensively. Therefore, we turned to a specialized group of commissural neurons, the precerebellar PN neurons in the hindbrain, whose cell bodies migrate tangentially over a considerable distance from the dorsal edge of the hindbrain to settle next to the ventral midline (Kawauchi et al., 2006; Okada et al., 2007; Shinohara et al., 2013). The migration of PN neurons towards the midline requires Netrin-1/ DCC/Robo3 signaling (Marillat et al., 2004; Yee et al., 1999). While the leading processes of PN neurons cross the midline, their cell bodies mostly terminate migration without midline crossing. This developmental feature enables considerable spatial separation of two PN populations: those in the early- and mid-migratory routes harboring the pre-crossing and those near the midline region harboring the post-crossing leading processes. We took advantage of this feature to obtain pure populations of pre- and post-crossing PN neurons and compared their transcriptomic profiles (results will be published elsewhere). We found that a pair of highly related class II bHLH TFs, *Nhlh1* and *Nhlh2*, were highly enriched in the pre-crossing (during migration), but were markedly down-regulated in the post-crossing (at final destination), PN neurons (enrichment: 54.52 fold for Nhlh1, 9.88 fold for Nhlh2). The differential expression of *Nhlh1* and *Nhlh2* was confirmed by in situ hybridization (ISH) expression data from the Allen Developing Mouse Brain Atlas (https://developingmouse.brain-map.org/) (Figure S1).

### Nhlh1 and Nhlh2 could induce Robo3 expression as transcription activators

The enrichment of Nhlh1 and Nhlh2 in pre-crossing PN neurons suggests that they might regulate the expression of genes required for the behavior of the pre-crossing PN neurons. Therefore, we screened for Nhlh1/2 binding sites in the promoters of transcripts that were enriched in the pre-crossing PN population from our RNA-seq data using the position frequency matrices (PFMs) of Nhlh1 and Nhlh2 from JASPAR. The screen revealed that the promoter of *Robo3*, a gene known to be highly expressed in migrating PN neurons (Marillat et al., 2004), bears multiple copies of potential Nhlh1/2 binding sites. Using the FIMO tool of the MEME Suite and the Eukaryotic Promoter Database, we scanned the proximal 2 kb promoter sequences of *Robo3*, and found seven statistically significant matches to the consensus Nhlh1/2 binding sequences (Figure 1A). Next, we aligned the promoter sequences of *Robo3* from six divergent mammalian species and identified a highly conserved region. The presence of the so-called phylogenetic footprints, though not direct proof, implies the potential importance of the conserved regions in the regulation of *Robo3* transcription. We found a highly conserved region within the 150 bp region proximal to the transcriptional start site (TSS) (Figure 1B) containing a Nhlh1/2 binding site with high matching probability (Figure 1A, C). The results from the in silico analysis raised the possibility that Nhlh1 and Nhlh2 might regulate *Robo3* transcription.

**Figure 1.**
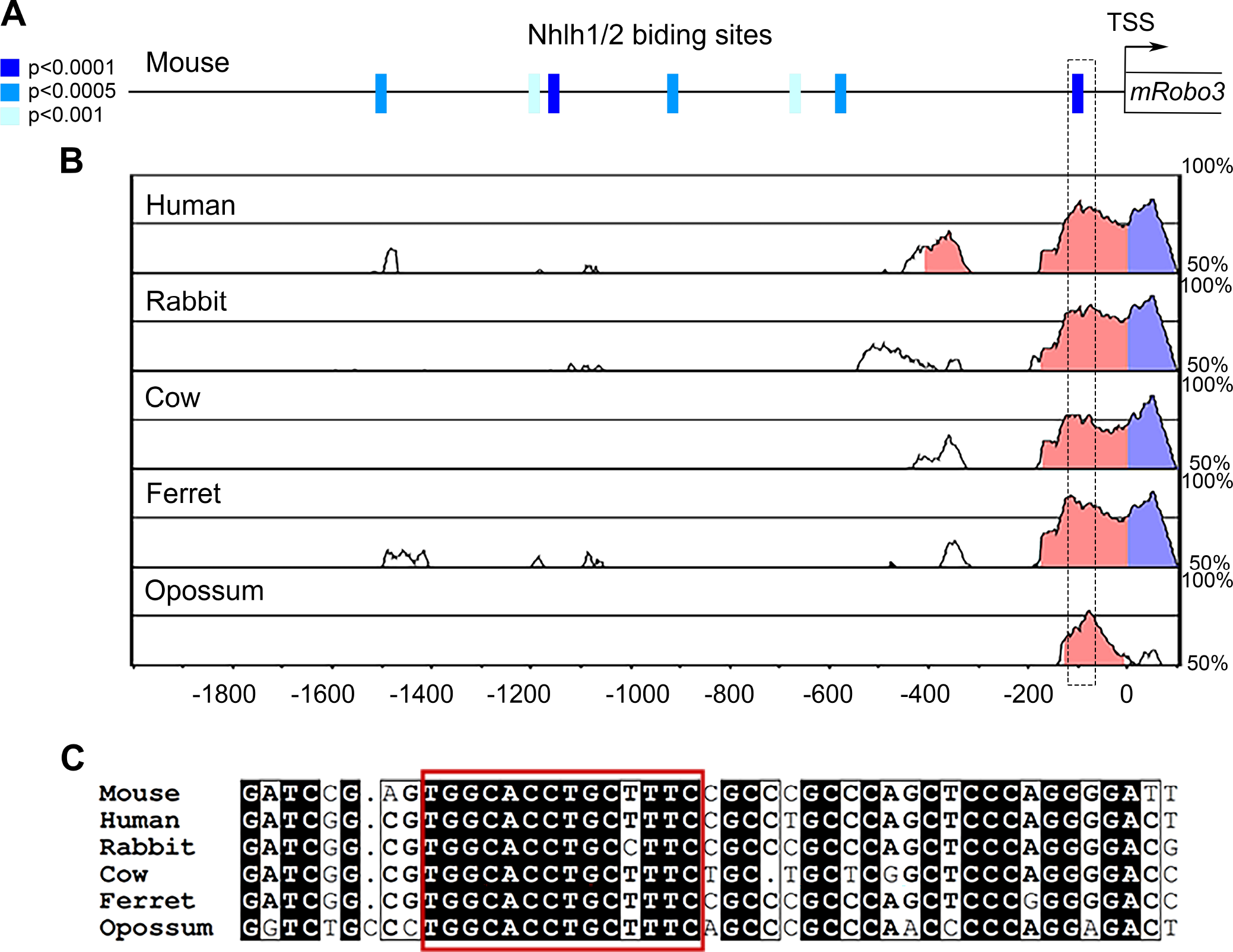
In silico analyses of Nhlh1/2 binding sites on *Robo3* promoter. (A) Predicted Nhlh1/2 binding sites across the 2 kb proximal promoter of mouse *Robo3* by FIMO (MEME Suites) are indicated by blue color-filled boxes. The shades of blue indicate the range of p-values as shown by the legend. TSS: transcriptional start site. (B) Highly conserved regions in the 2 kb proximal promoters of *Robo3* across six divergent mammalian species were detected using mVISTA. The Y axes indicate percentage of identity. Pink and blue colored areas indicate regions with peak homology values above 70% upstream and downstream, respectively, of TSS. (C) Sequence alignment of the highly conserved regions indicated by the box with dashed line in (A) & (B) in the Robo3 promoters across six mammalian species. The red box in (C) highlights the sequences of the predicted Nhlh1/2 binding site.

We then tested whether forced expression of Nhlh1 and Nhlh2 could induce ectopic Robo3 expression. We introduced expression vectors of full-length *Nhlh1* and *Nhlh2* into the lower rhombic lip of the E12.5 mouse hindbrains via in utero electroporation (EP) and analyzed Robo3 protein expression at E14.5 (Figure 2A). The lower rhombic lip region situated at the dorsal edge of the caudal hindbrain (Figure 2B) contains the progenitor zone of PN neurons (Altman and Bayer, 1987; Pierce, 1966; Rodriguez and Dymecki, 2000; Wingate and Hatten, 1999), but is devoid of Robo3 expression (Figure 2C non-EP side, 2E) because post-mitotic PN neurons only begin to express Robo3 after leaving the rhombic lip. Forced expression of Nhlh1 and Nhlh2 in the rhombic lip clearly induced the ectopic expression of Robo3 within the rhombic lip region (Figure 2C, C’, C’’, D, D’, D’’). Electroporated neurons found deep within the hindbrain neuroepithelium also expressed ectopic Robo3 (Figure 2F, F’, F’’). Ectopic Robo3 induction was also observed when *Nhlh1* and *Nhlh2* were electroporated into the midbrain (Figure 2G, H, H’, I, I’, I’’ compared to J). EP of *Nhlh1* or *Nhlh2* alone also induced ectopic Robo3 expression in the rhombic lip and midbrain, although it appeared to be less efficient than EP of both *Nhlh1* and *Nhlh2* (Figure S2A, B). In contrast, EP of full-length *Lhx2*, a molecule previously shown to directly control *Robo3* transcription in dI1 spinal neurons (Kawauchi et al., 2010; Wilson et al., 2008), did not induce ectopic Robo3 expression in the rhombic lip and midbrain (Figure S2C). *Robo3* ISH on EP samples indicated that the ectopic expression of *Robo3* was induced at the transcription level (Figure S2D). Taken together, these results suggest that Nhlh1 and Nhlh2 regulate *Robo3* transcription.

**Figure 2.**
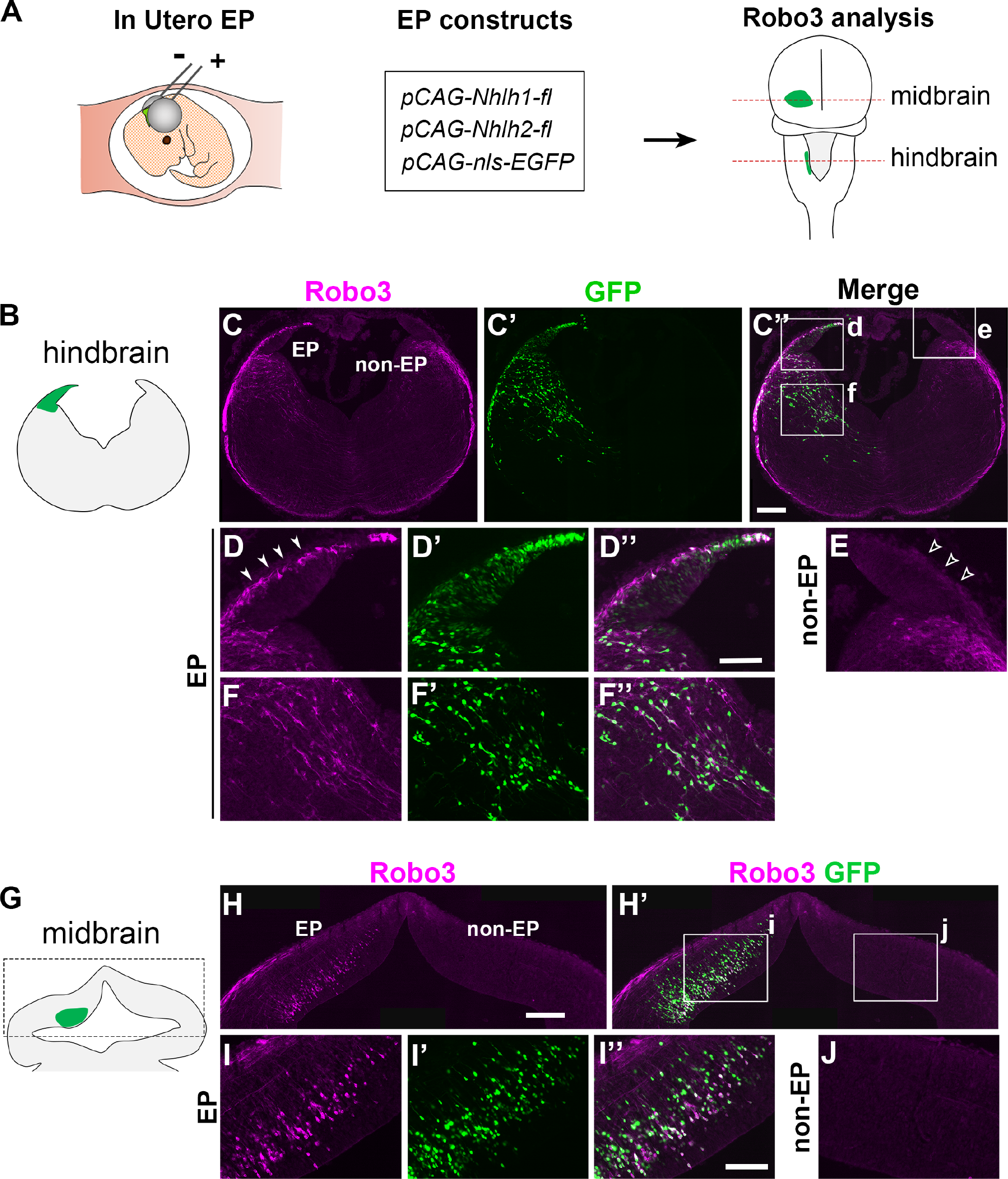
Forced expression of full-length Nhlh1 and Nhlh2 induce ectopic Robo3 expression in the hindbrain and midbrain. (A) A schematic showing the experimental procedure. DNA constructs were introduced into the hindbrain or midbrain via in utero EP at E12.5, and the EP samples were analyzed at E14.5 on sections by Robo3 and GFP IHC. The approximate positions of sections shown for each region are indicated by red dashed lines. (B) A schematic showing a hindbrain section with EP unilaterally targeted to the dorsal hindbrain including the rhombic lip region (green). (C), (C’) & (C’’) A hindbrain section at low magnification with Robo3 and GFP IHC. EP and non-EP stand for electroporated and non-electroporated sides, respectively. The EP side, but not the non-EP side, showed induction of ectopic Robo3 expression (n = 6). (D), (D’) & (D’’) High magnification images of EP area (d) marked in (C’’). The filled arrowheads indicate ectopic expression of Robo3 within the rhombic lip region. (E) A high magnification image of the non-EP area (e) marked in (C’’). The hollow arrowheads indicate lack of Robo3 induction within the rhombic lip. (F), (F’) & (F’’) High magnification images of the EP area (f) in (C’’). Many GFP positive neurons deep in the hindbrain epithelium also express ectopic Robo3. (G) A schematic showing a midbrain section with EP unilaterally targeted to the dorsal midbrain (green). The boxed area shows the approximate region shown in (H) & (H’). (H) & (H’) A midbrain section at low magnification with Robo3 and GFP IHC. The EP side, but not the non-EP side, showed induction of ectopic Robo3 expression (n = 3). (I), (I’), (I’’) High magnification images of the EP area (i) marked in (H’). (J) A high magnification image of the non-EP area (j) marked in (H’). Scale bars: 200 μm in (C), (C’) & (C’’); 100 μm in (D), (D’), (D’’), (E), (F), (F’) & (F’’); 200 μm in (H) & (H’); 100 μm in (I), (I’), (I’’) & (J).

Next, we asked whether Nhlh1 and Nhlh2 function as transcriptional activator or repressor, and which domains of these molecules are important for the induction of Robo3. Nhlh1 and Nhlh2 share a highly homologous canonical bHLH domain in the C-terminus and a poorly conserved low-complexity domain in the N-terminus (Figure 3A) (Begley et al., 1992; Brown et al., 1992). In between the two domains, both molecules contain a highly conserved stretch of 11 amino acids, encompassing six or five consecutive arginine residues, immediately preceding the canonical bHLH domain. This stretch, which we named the R6 domain, is a feature specific to Nhlh1 and Nhlh2 (Begley et al., 1992; Brown et al., 1992). The highly charged R6 domain may form an extended basic domain together with the canonical basic region, rendering the DNA binding specificity unique to Nhlh1/2 (Brown and Baer, 1994). Evidence exists on both sides for the role of Nhlh1/2 as an activator or repressor (Good and Braun, 2013; Isogai et al., 2011; Kruger et al., 2004; Manetopoulos et al., 2003; Wankhade and Good, 2011). If Nhlh1/2 induce Robo3 expression by directly binding to its promoter, as suggested by our in silico analysis, it would imply that Nhlh1/2 works as transcriptional activators. To test this possibility, we generated two types of Nhlh1/2 fusion molecules: (1) fusion with a potent trans-activating domain from the herpes simplex virus TF VP16 (Sadowski et al., 1988) and (2) fusion with a potent repressor domain, EnR, from the Drosophila Engrailed (Fan and Sokol, 1997) (Figure 3B). We electroporated the fusion constructs into E12.5 brainstems as in Figure 2 and assayed for Robo3 induction in the rhombic lip and the midbrain. VP16-fused full length Nhlh1/2 strongly induced Robo3 expression in the rhombic lip (Figure 3E) as well as in the midbrain (Figure S2E). In contrast, EnR-fused Nhlh1/2 failed to induce ectopic Robo3 expression in the rhombic lip (Figure 3F) and midbrain (Figure S2F). These results suggest that Nhlh1/2 serve as transcriptional activators to induce Robo3 expression. Next, we investigated the functional importance of the R6 domain in Robo3 induction. We found that VP16 fused canonical bHLH domain without the R6 domain did not induce Robo3 expression (Figure 3G, Figure S2G), whereas VP16 fused R6-containing bHLH domain was a potent inducer of Robo3 (Figure 3H, Figure S2H). These results suggest that the R6 domain unique to the Nhlh1/2 subfamily, together with the canonical bHLH domain, is essential for Nhlh1/2 to activate the expression of Robo3.

**Figure 3.**
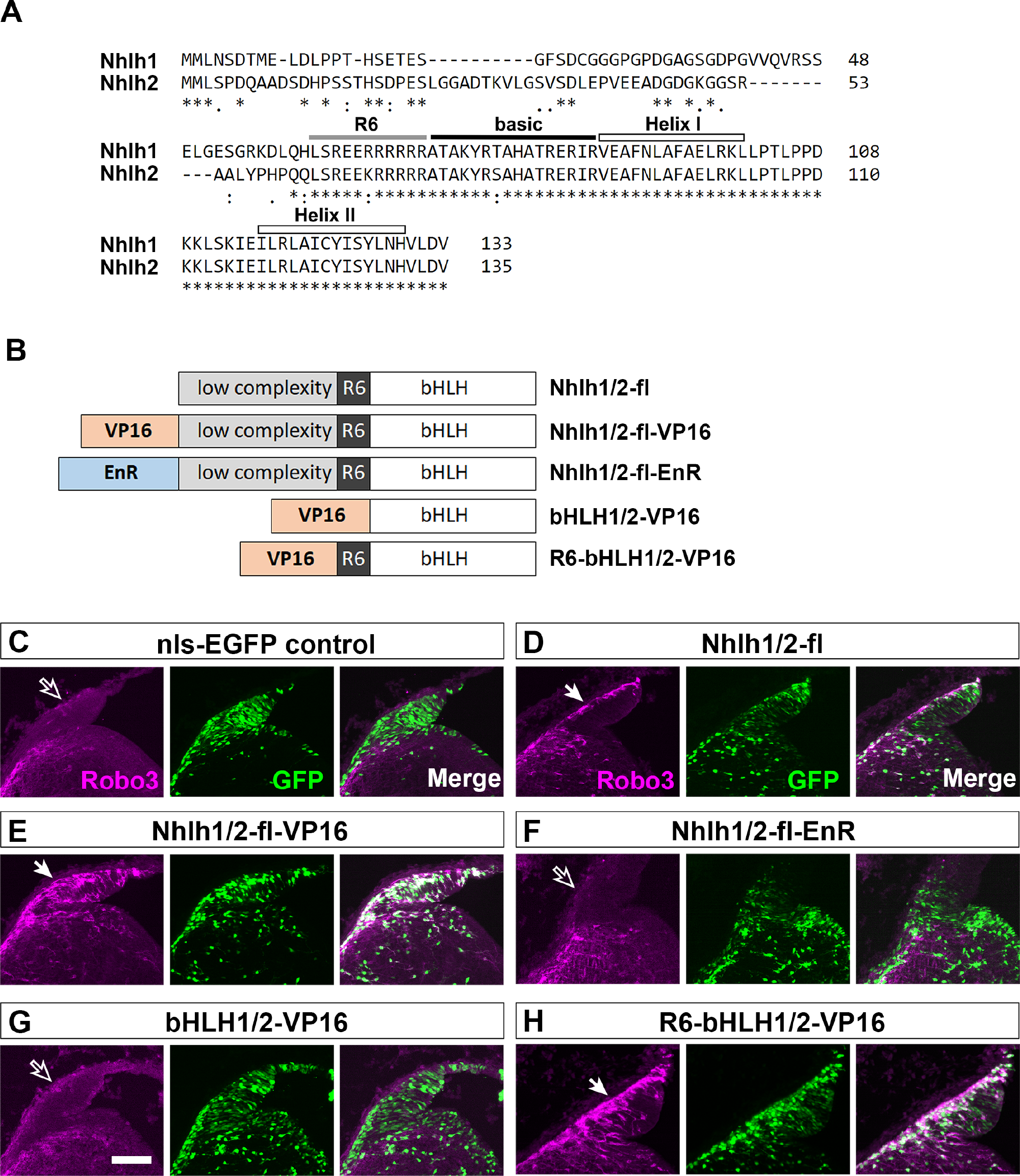
Nhlh1 and Nhlh2 are transcriptional activators in Robo3 induction and the arginine-rich domain upstream of bHLH domain is important for their inducibility. (A) Sequence alignment of Nhlh1 and Nhlh2 proteins shows a highly conserved C-terminal half encompassing a canonical bHLH domain and a stretch of 11 amino acids upstream of the bHLH with a consecutive run of arginine (R) (coined R6 domain in this study). The N-terminal halves of the two proteins are far less conservative and of low complexity structurally. (B) A schematic of constructs for testing Robo3 inducibility. VP16 is the potent transcriptional activation domain from the herpes simplex virus protein VP16. EnR is the potent transcriptional repressor domain from the drosophila protein Engrailed. (C), (D), (E), (F), (G) & (H) The rhombic lip regions of hindbrains electroporated with the indicated DNA constructs. Filled arrows indicate ectopic Robo3 induction in the rhombic lip in (D, n= 6), (E, n =3) and (H, n = 3), while hollow arrows indicate lack of Robo3 induction in (C, n = 5), (F, n = 3) and (G, n = 3). Scale bar: 100 μm in (C), (D), (E), (F), (G) & (H).

### Forced expression of Nhlh1/2 drives contralateral axon projections

Since forced Nhlh1 and Nhlh2 expression induce ectopic Robo3 expression, we wondered if it could instruct axon projections towards the ventral midline? To address this, we force-expressed Nhlh1 and Nhlh2 in the developing hindbrain and midbrain as in Figure 2, and examined the axon trajectories of the electroporated neurons. In the midbrain, we employed a previously established method that enabled the examination of axon laterality in flat-mounted brainstem after in utero EP (Figure 4A) (Inamata and Shirasaki, 2014). EP with an *mCherry* expression construct into the midbrain at E11.5 labelled mostly caudally-extending ipsilateral axons, in line with the previous report (Figure 4B, C) (Inamata and Shirasaki, 2014). Forced expression of Nhlh1 and Nhlh2 with mCherry at E11.5 could direct some of these neurons to project contralaterally (Figure 4B, C). In the hindbrain, control EP with a *GFP* construct at E12.5 labelled only a few neurons whose axons extend ventrally towards midline, resulting in low levels of GFP fluorescence near the midline region (Figure 4D, E). In contrast, EP of *Nhlh1* and *Nhlh2* with *GFP* labelled numerous neurons that either extended their axons or migrated towards the midline (Figure 4D, E). The GFP signal near the midline was significantly higher than that of the control (Figure 4E). These results from both the midbrain and hindbrain show that forced expression of Nhlh1/2 could drive changes in axonal projection from ipsilateral to contralateral, suggesting that axon laterality in vivo could be determined by the presence or absence of a pair of TFs.

**Figure 4.**
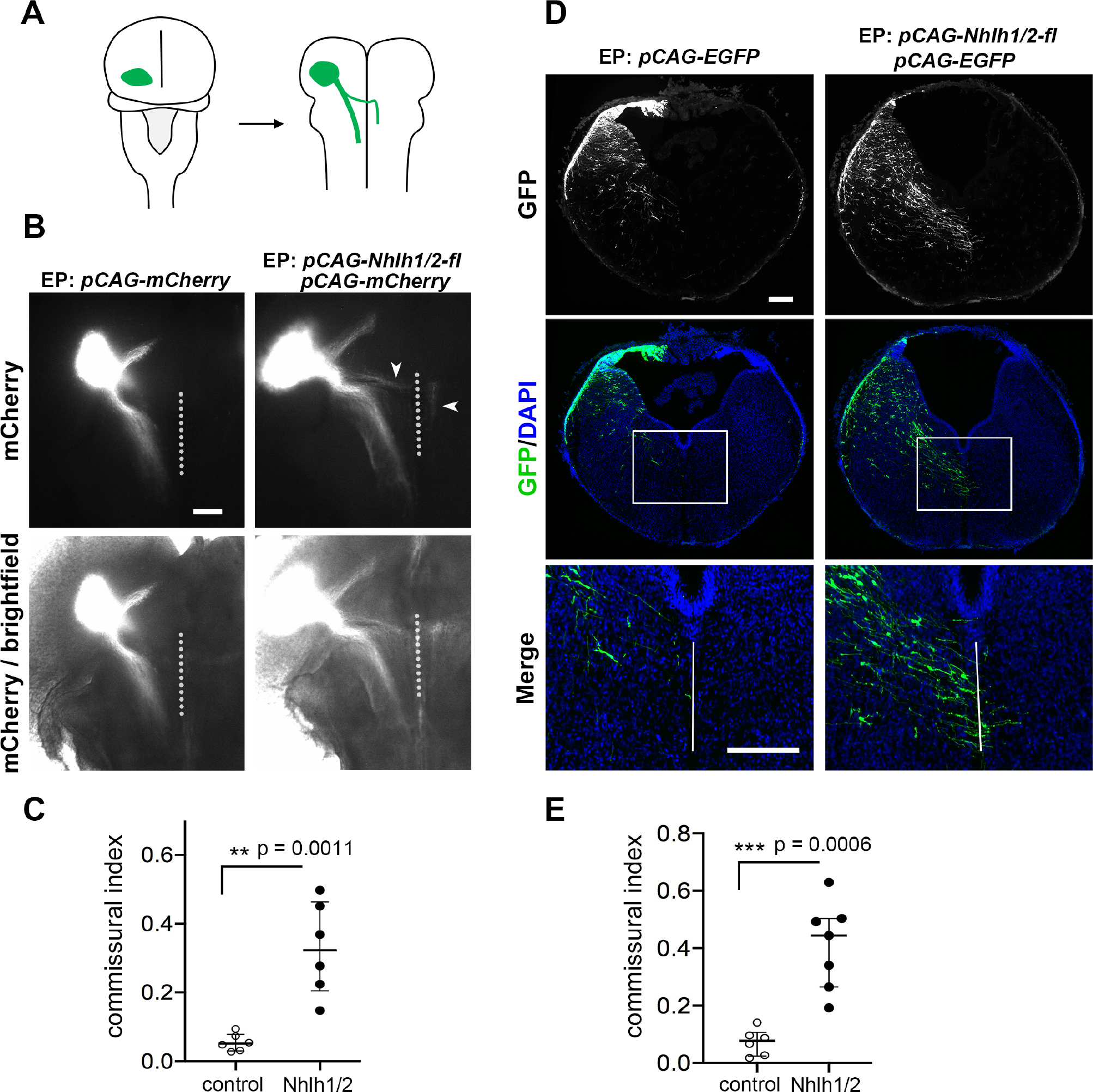
Forced expression of full-length Nhlh1 and Nhlh2 drive axonal projection towards the ventral midline. (A) A schematic showing the experimental procedure of testing the effect of force-expressing Nhlh1/2 on midbrain axonal trajectories. EP was performed at E11.5 into the dorsal midbrain, and the analysis of axonal trajectories of EP neurons was carried out on flat-mounted brainstems at E14.5. (B) Partial views of samples electroporated with either *mCherry* as control (n = 6), or with full-length *Nhlh1/2* and *mCherry* constructs (n = 6). Fluorescence images were shown in the top panel, and those that merged with the brightfield images were shown in the bottom panel to visualize the positions of the ventral midline (indicated by vertical lines). (C) Quantification of the midbrain EP experiment. The commissural index is a measurement of the relative quantity of axons projecting to and reaching the ventral midline over those that projecting ipsilaterally (see materials and methods). Data were represented by a scatter plot, with the median and the upper and lower quantile indicated. Force-expressing Nhlh1 and Nhlh2 increased significantly the proportion of axons that crossed the ventral midline (p = 0.0011, Mann-Whitney U test). (D) The effect of expressing full-length *Nhlh1/2* (n = 6) in comparison to expressing only *EGFP* construct (n = 7) on the trajectories of hindbrain axons. EP was performed at E12.5 into the dorsal hindbrain, and the analysis of axonal trajectories was carried out at E14.5 on sections of EP samples after GFP IHC. The vertical lines indicate the ventral midlines, and the bottom panel shows the high magnification images around the ventral midline regions corresponding to the boxed areas in the middle panel. (E) The quantification of the hindbrain EP experiment. The commissural index reflects the relative quantity of both axons and neurons reaching the vicinity of the ventral midline over those that are immediately below the rhombic lip (see materials and methods). Data were represented by a scatter plot, with the median and the upper and lower quantile indicated. Force-expressing Nhlh1 and Nhlh2 increased significantly the proportion of axons and neurons that reached the ventral midline (p = 0.0006, Mann-Whitney U test). Scale bars: 400 μm in (B); 200 μm in (D).

### Expression of Nhlh1 and Nhlh2 in relation to Robo3 during the development of commissural neurons

Co-localization of Nhlh1 and Nhlh2 with Robo3 should be a strong indicator of whether and where these molecules may regulate Robo3 transcription in the CNS. Robo3 has previously been shown to be expressed and required in almost all commissural neurons whose axons cross the ventral midline at the FP, which spans from the spinal cord to the midbrain (Friocourt et al., 2019; Inamata and Shirasaki, 2014; Marillat et al., 2004; Sabatier et al., 2004; Tamada et al., 2008). In the forebrain, the FP ceases to exist (Placzek and Briscoe, 2005), and Robo3 expression is absent from the forebrain commissural tracts (Friocourt et al., 2019). We examined the expression patterns of Nhlh1 and Nhlh2 in relation to that of Robo3 in areas and at stages where and when FP-crossing commissural axon projections take place.

Because good antibodies to detect or distinguish Nhlh1 and Nhlh2 are unavailable, we turned to ISH using *Nhlh1*, *Nhlh2*, and *Robo3* riboprobes on adjacent sections. *Nhlh1*, *Nhlh2* and *Robo3* expression was first examined in migrating PN neurons at E14.5. Consistent with our RNA-seq and the ISH expression data from the Allen Developing Mouse Brain Atlas (https://developingmouse.brain-map.org/) (Figure S1), we found that both *Nhlh1* and *Nhlh2* are expressed in migrating PN neurons, similar to *Robo3* expression (Figure 5A). Next, we examined the expression of these three molecules across the entire neuroaxis of the CNS at E11.5 and E13.5, which are stages when most commissural neurons project axons towards the ventral midline. We found that *Robo3* generally labelled neurons immediately adjacent to the ventricular zone, where young immature neurons reside, but not more differentiated neurons in superficial positions (Figure 5B-H). This is consistent with previous findings that Robo3 is transiently expressed in pre-crossing commissural neurons and is downregulated soon after commissural axons cross the midline (Marillat et al., 2004; Sabatier et al., 2004).

**Figure 5.**
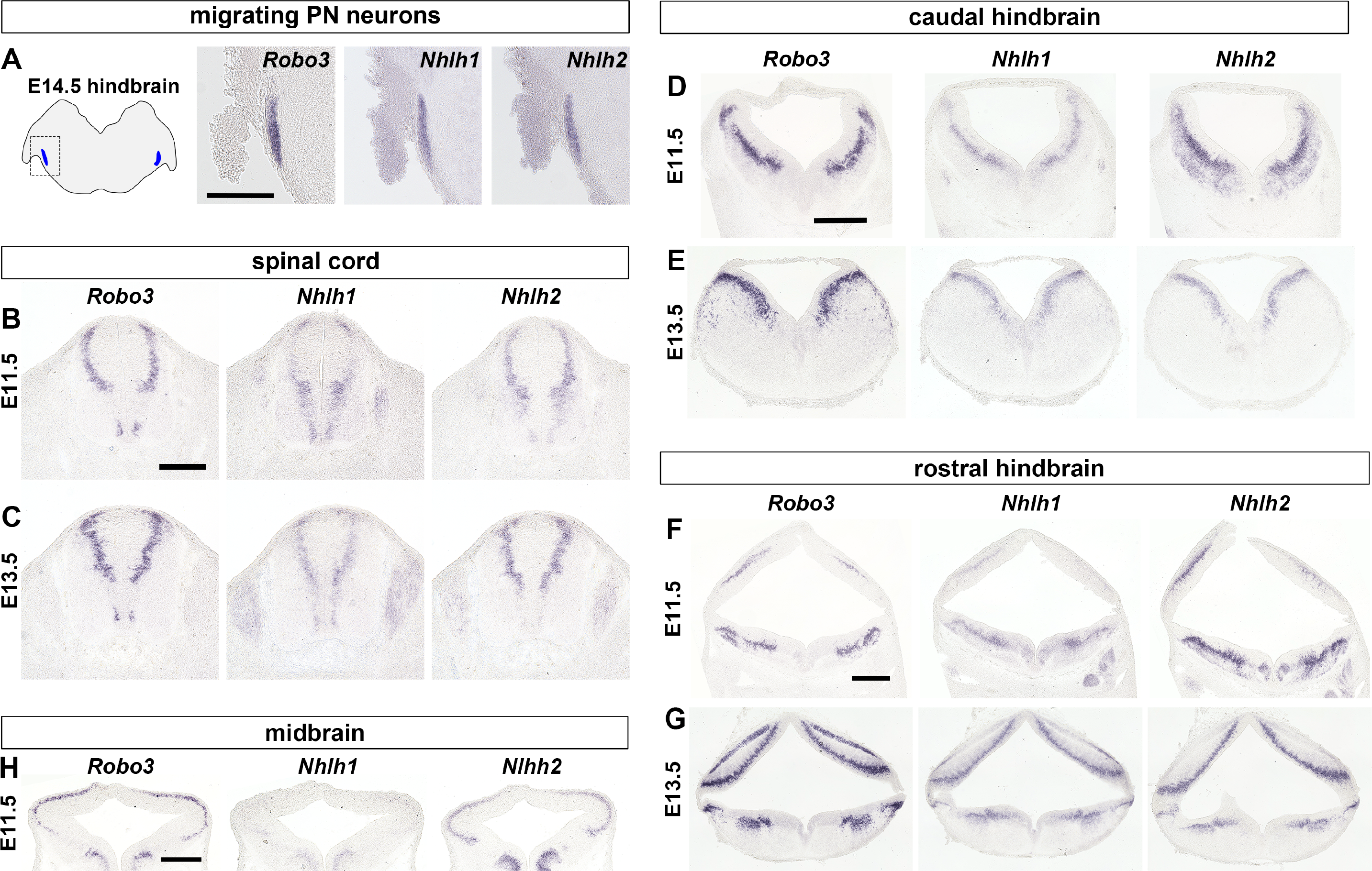
Expression of *Robo3*, *Nhlh1* and *Nhlh2* on adjacent sections from the spinal cord to the midbrain by ISH. (A) The schematic on the left shows a hindbrain section at E14.5 with the cross-sections of migrating PN neurons in blue. The boxed area indicates region of the ISH images shown on the right. (B) & (C) Spinal cord sections at E11.5 and E13.5, respectively. (D) & (E) Caudal hindbrain sections at E11.5 and E13.5, respectively. (F) & (G) Rostral hindbrain sections at E11.5 and E13.5, respectively. (H) Midbrain sections at E11.5. Note the E13.5 midbrain data is not shown as *Robo3* expression in the midbrain at this stage is very weak. Scale bars: 200 μm in (A); 200 μm in (B) & (C); 400 μm in (D) & (E); 400 μm in (F) & (G); 200 μm in (H).

Remarkably, from the spinal cord to the midbrain, *Nhlh1* and *Nhlh2* were expressed in areas that also expressed *Robo3* (Figure 5B- H). Two points should be noted here. First, combined expression of *Nhlh1* and *Nhlh2* appears wider than that of *Robo3*, which is more obvious at E11.5 than E13.5. For example, *Nhlh1* and *Nhlh2* seem to be expressed, albeit weakly, in the ventral spinal cord which is largely devoid of *Robo3* (Figure 5B).

Likewise, in the E11.5 caudal and rostral hindbrains (Figure 5D, F), *Nhlh2*, in particular, shows expression in more differentiated neurons located superficially. Second, the relative strength of *Nhlh1* and *Nhlh2* expression appears to vary depending on the brain region and neuronal subdomain. For example, *Nhlh1* expression appears to be stronger than *Nhlh2* expression in the ventral-most spinal domain, whereas *Nhlh2* appears stronger in the dorsal spinal cord (Figure 5B). Despite the varying relative level of expression, the *Nhlh1* and *Nhlh2* expression domains mostly overlap. In the E11.5 midbrain, the *Nhlh1* signal is very weak but is still present as a faint band of signal correlating to the *Robo3* pattern. The results from the expression study of *Nhlh1*, *Nhlh2* and *Robo3* suggest the following: (i) a high likelihood that Nhlh1 and Nhlh2 may regulate Robo3 expression in vivo in most commissural axons that cross the midline via the FP; and (ii) Nhlh1 and Nhlh2 may functionally complement each other. However, in brain regions rostral to the midbrain, *Robo3* and *Nhlh1* and *Nhlh2* expression was much less correlated (Figure S6A, B, C).

### FP-crossing commissural axons from the spinal cord to the midbrain fail to approach the ventral midline in *Nhlh1* and *Nhlh2* double deficient mice

To examine the endogenous function of Nhlh1 and Nhlh2, we generated *Nhlh1* and *Nhlh2* deficient mice by CRISPR/Cas9 mediated gene editing in germ line cells (Figure S3A, B). The resultant *Nhlh1* mutant allele (*Nhlh1-m*) carries a premature termination site just before the first helix region, thus generating a truncated Nhlh1 protein without the HLH region (Figure S3C). The *Nhlh2-m* allele carries corrupted amino acids over the extended basic domain and the first helix regions (Figure S3D), resulting in a peptide that could no longer generate the bHLH structure. Given that the bHLH region is essential for DNA binding and protein-protein interactions of bHLH TFs (Bertrand et al., 2002; Dennis et al., 2019), we expected that *Nhlh1-m* and *Nhlh2-m* are likely to be loss-of-function alleles. To confirm this, we cloned the coding sequences of *Nhlh1-m* and *Nhlh2-m* into expression vectors. Forced expression of Nhlh1-m and Nhlh2-m in E12.5 mouse embryos by EP as in Figure 2 did not induce ectopic expression of Robo3 in the rhombic lip (Figure S3E). We also confirmed that Nhlh1-m and Nhlh2-m did not act dominant negatively in suppressing Robo3 expression, as PN neurons expressing Nhlh1-m and Nhlh2-m showed normal Robo3 expression (Figure 3F).

We then generated single and double homozygotes of *Nhlh1-m* and *Nhlh2-m*. Targeted knockout of *Nhlh1* and *Nhlh2* has been previously generated (Cogliati et al., 2002; Good et al., 1997; Kruger and Braun, 2002) and in addition to perinatal lethality, only two defects in the developing brain have been reported in the double knockout (dKO) mice: defects in the migration of PN neurons (Schmid et al., 2007) and Gonadotropin releasing hormone expressing neurons (GnRH) (Kruger et al., 2004). However, the mechanisms underlying these defects have not been elucidated. We first analyzed PN formation in the double *Nhlh1/2* mutant we generated. We found that PN neurons failed to migrate towards the ventral midline, but instead arrested migration in the lateral and anteriorly-extended positions (Figure 6A, B). This phenotype observed in our mice resembled that previously reported in dKO mice (Schmid et al., 2007). PN formation in single *Nhlh1* and *Nhlh2* mutants appeared normal, again in line with previous reports (Figure S4A) (Schmid et al., 2007). Interestingly, the PN phenotype observed in the present study and in the previous dKO is highly reminiscent of the PN defect in the *Robo3* mutant (Marillat et al., 2004). Another precerebellar nucleus, the inferior olivary nucleus (ION), is also affected in the *Robo3* mutant (Di Meglio et al., 2008; Marillat et al., 2004). Therefore, we examined ION in the *Nhlh1/2* double mutant. We found that in the double mutant, ION neurons failed to gather tightly around the ventral midline as in the control, but were situated at a small distance away from the midline, a phenotype that again resembled that of the *Robo3* mutant (Figure S4B, C) (Di Meglio et al., 2008).

**Figure 6.**
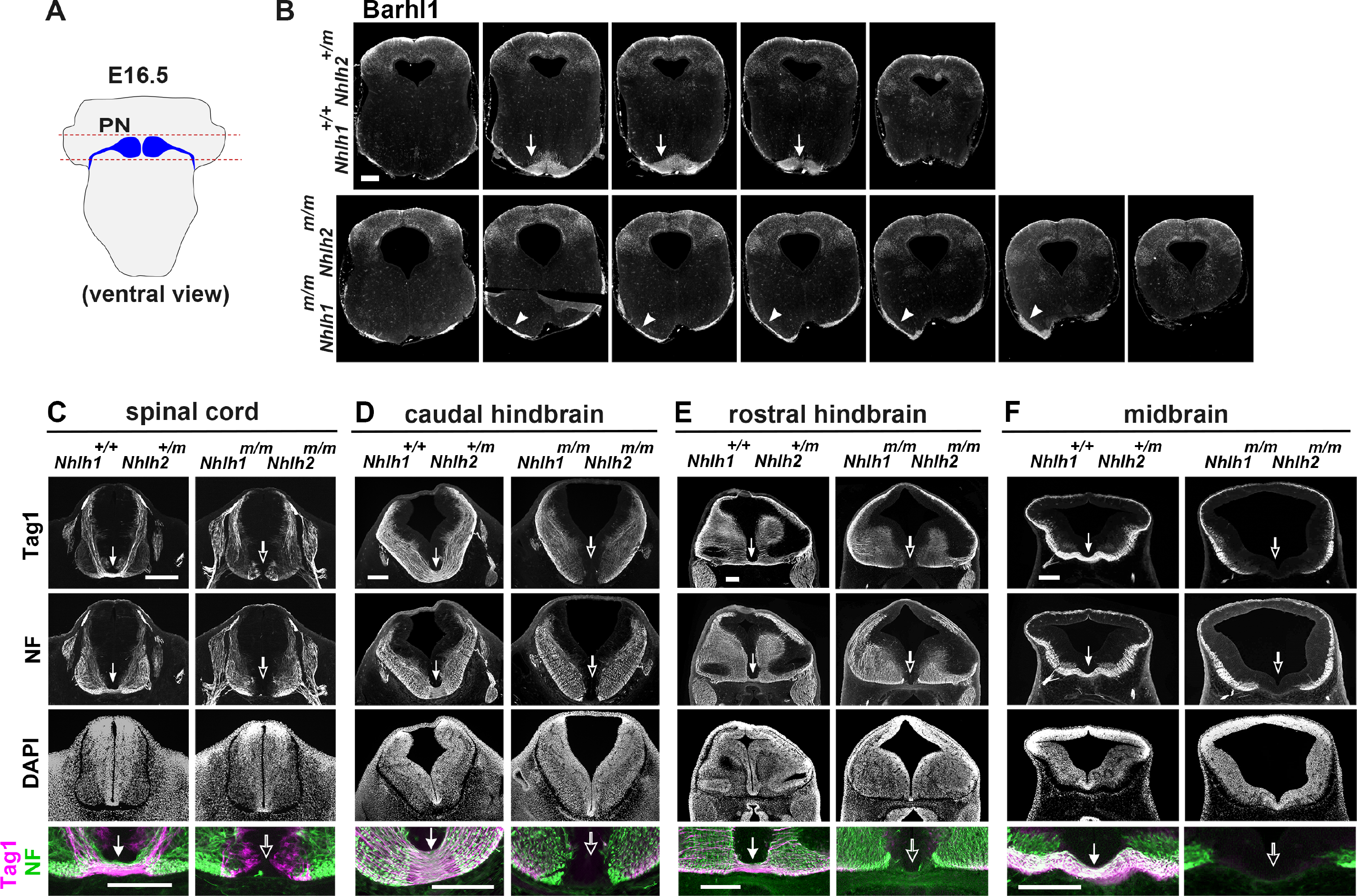
Lateralized PN and lack of ventral commissures in the spinal cord, hindbrain and midbrain in *Nhlh1* and *Nhlh2* double mutant. (A) A schematic showing the PN neuron migration and PN formation in an E16.5 hindbrain. Red dashed lines mark the rostrocaudal span of the sections shown in (B). (B) Sections across the PN region from an E16.5 hindbrain of the control genotype (top panel) and the double mutant (lower panel) with PN marked by Barhl1 IHC. PN neurons form a nucleus next to the ventral midline (arrows) (n = 2) in the control genotype, but were positioned in lateralized positions (arrowheads) indicating failure to approach the ventral midline (n = 2). (C), (D), (E) & (F) coronal sections from the spinal cord (C), caudal hindbrain (D), rostral hindbrain (E), and midbrain (F) at E11.5 were subjected to Tag1 and NF double IHC. The Tag1 signals label the commissural axons and the NF signals depict the general axonal patterns. The DAPI counterstaining indicate the overall cytoarchitecture. Comparisons were made between the control genotype (*Nhlh1^+/+^ Nhlh2^+/m^*) (n = 3) and the double mutant (*Nhlh1^m/m^ Nhlh2^m/m^*) (n = 3). The Tag1 and NF merged images in the bottom panel are high magnification images of the ventral commissure regions. Filled arrows show the ventral commissures and the hollow arrows show the lack of ventral commissures. Scale bars: 200 μm in (B); 200 μm in low magnification images in (C), (D), (E) & (F); 200 μm in high magnification images in (C), (D), (E) & (F).

The grossly correlated expression of *Nhlh1*, *Nhlh2* and *Robo3* in commissural neurons from the spinal cord to the midbrain prompted us to examine ventral commissure formation throughout these regions in our mutant mice. Tag1 (also known as Contactin2) was used as a marker for commissural axons (Dodd et al., 1988; Yamamoto et al., 1986), and neurofilament (NF) immunohistochemistry (IHC) was used to reveal the overall axonal patterns. We found that ventral commissures completely failed to form in the double mutant in the spinal cord (Figure 6C), caudal hindbrain (Figure 6D), rostral hindbrain (Figure 6E) and midbrain (Figure 6F). The overall axonal patterns, except for the lack of ventral commissures and the cytoarchitecture of these CNS regions shown by DAPI staining, appeared grossly normal (Figure 6C-F). Tag1 positive axons initially developed normally but failed to converge and extend all the way to the ventral midline, which was better appreciated in the spinal cord samples (Figure 6C). No notable abnormalities in the ventral commissure formation were observed in the single mutants of *Nhlh1* and *Nhlh2* (Figure S4D-G). The prevalent absence of ventral commissures persisted into later stages, as shown by analyses of the E13.5 and E16.5 spinal cord (Figure S4H, I), and E13.5 hindbrain and midbrain (Figure S4J, K).

What about the non-FP crossing commissural axons, such as dorsally crossing commissural axons or commissures formed anterior to the midbrain where the FP ceases to exist, whose extension to the midline has been shown previously to be independent of Robo3 (Chedotal, 2014; Comer et al., 2015; Friocourt et al., 2019)? We investigated whether these other types of commissural axons could form normally in our *Nhlh1/2* double mutant. We found that the dorsal commissure in the spinal cord, anterior commissure in the basal forebrain and corpus callosum were all normally formed in the double mutant (Figure S6D, E, F). This result indicates that Nhlh1/2 deficiency specifically affects commissural axons that cross the midline via the FP.

### Marked reduction of Robo3 in *Nhlh1* and *Nhlh2* double mutant from the spinal cord to the midbrain

We showed that FP-crossing commissures failed to form in the *Nhlh1/2* double mutant, resembling the defects observed in the *Robo3* mutant. Therefore, we investigated Robo3 expression in the *Nhlh1/2* double mutant by Robo3 IHC. We found a huge reduction in Robo3 expression in the E11.5 spinal cord (Figure 7A), caudal hindbrain (Figure 7B), rostral hindbrain (Figure 7C), and midbrain (Figure 7D) of the double mutant compared to the control. Similarly, a marked reduction in Robo3 was also observed in the migrating PN neurons at E16.5 (Figure 7E). Examination of *Robo3* transcription by ISH also revealed a huge reduction in *Robo3* mRNA in the spinal cord (Figure 7F), and the migrating PN neurons (Figure 7G) in the double mutant. These results provide the ultimate proof that Nhlh1 and Nhlh2 transactivate Robo3 expression in vivo.

**Figure 7.**
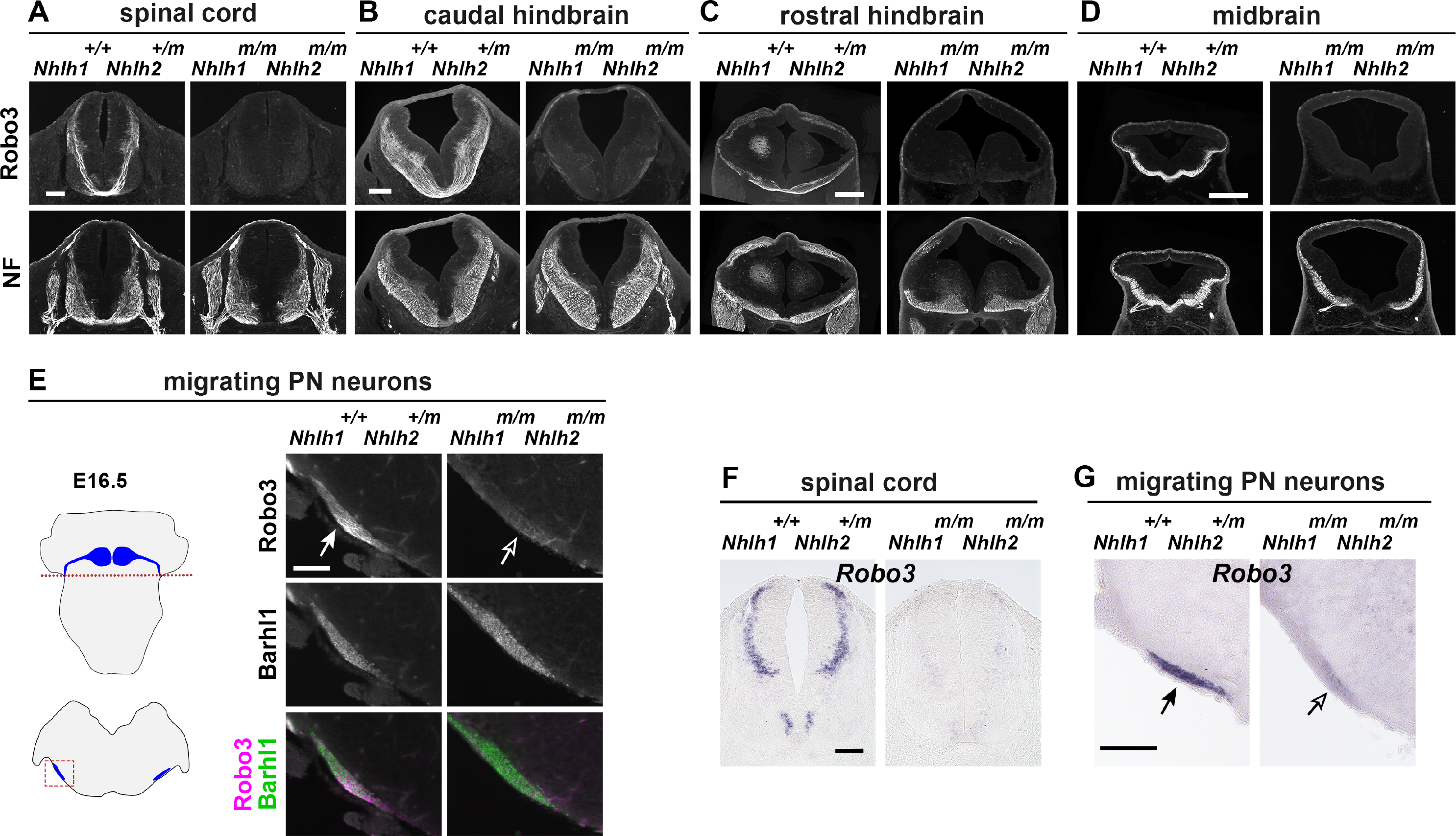
Huge reduction of Robo3 expression in commissural neurons in *Nhlh1* and *Nhlh2* double mutant. Coronal sections from the spinal cord (A), caudal hindbrain (B), rostral hindbrain (C), and midbrain (D) at E11.5 were subjected to Robo3 and NF double IHC. A huge reduction of Robo3 expression was detected in the double mutant (n = 3) in comparison with the control genotype (n = 3) at these axial levels. (E) The schematics on the left show the migrating PN neurons in a whole mount E16.5 hindbrain in the top, and on a coronal section in the bottom. The red dashed line and the red boxed area indicate the approximate axial level and the area, respectively, of IHC images on the right. PN neurons were labelled by Barhl1 IHC. A huge reduction of Robo3 expression was detected in the PN neurons in the double mutant (hollow arrow) (n = 2) in comparison with the control genotype (arrow) (n = 2). (F) & (G) *Robo3* ISH on E11.5 spinal cord and E16.5 hindbrain sections, respectively, showed that *Robo3* mRNA was markedly reduced in the double mutant. Scale bars: 100 μm in (A); 200 μm in (B); 400 μm in (C) & (D); 100 μm in (E); 100 μm in (F); 200 μm in (G).

We believe that Robo3 deficiency is the main cause of the commissure-less phenotype for the following reasons. First, Robo3 was barely detectable in the Nhlh1/2 double mutant. Second, we examined the expression of *Netrin-1* and *Shh*, the two main guidance molecules previously shown to guide commissural axons towards the midline (Charron et al., 2003; Kennedy et al., 1994; Serafini et al., 1996), and found that they were expressed in a similar manner in the double mutant and the control (Figure S5A, B). Third, neurogenesis and initial axon extension of commissural neurons did not appear to be affected in the double mutants. This is because Tag1 and DCC, both predominantly expressed in commissural neurons, showed almost normal expression profiles in the dorsal half of the spinal cord in the double mutant where most commissural neurons reside (Figure 6C, Figure S5C). In addition, Brn3a+ neurons, which are dI1, dI2, dI3, dI5, and dILB spinal neuronal classes comprising a mixture of ipsilateral- and contralateral-projecting neurons (Lai et al., 2016; Tulloch et al., 2019), were comparable between the double mutant and the control (Figure S5D). Similarly, the specification of the PN neurons also appeared to be normal in the double mutant, as these neurons migrated anteriorly and expressed the PN neuronal markers Barhl1 and DCC (Figure 6B, Figure S5E).

### Robo3 expression rostral to the midbrain is not affected in Nhlh1/2 double mutant

The seemingly uncorrelated expression of *Nhlh1*, *Nhlh2*, and *Robo3* in structures rostral to the midbrain (Figure S6A) raised the possibility that Robo3 expression in these structures might not be affected in the *Nhlh1/2* double mutant. To test this prediction, we examined Robo3 expression in three forebrain structures known to express Robo3: the ganglionic eminence (GE), hypothalamus (HTH), and medial habenular nucleus (mHb) (Figure S6A) (Barber et al., 2009; Belle et al., 2014; Quina et al., 2009; Schmidt et al., 2014). We found no differences in Robo3 expression levels in the GE, HTH and mHb between the double mutant and control (Figure S6G, H, I). Axons from the mHb form a highly fasciculated fasciculus retroflexus (FR) that projects caudally through the midbrain and crosses the ventral midline at the midbrain/hindbrain junction (Belle et al., 2014). We found that the FR in the double mutant continued to express Robo3 (Figure S6J) and was able to approach and cross the ventral midline (Figure S6K), supporting our earlier conclusion that the extrinsic guidance program for midline crossing is intact in the double mutant. Taken together, these results suggest that Robo3 expression in the forebrain is regulated by transcriptional program other than Nhlh1/2.

## DISCUSSION

In this study, we uncovered a transcriptional program involving the bHLH TFs, Nhlh1 and Nhlh2, for regulating the contralateral projection of commissural axons in mice. We initially identified these molecules in a specialized commissural system, the hindbrain PN neurons. Remarkably, mice deficient in both molecules showed that Nhlh1 and Nhlh2 have a global function in all FP-crossing commissural neurons from the spinal cord to the midbrain. We further showed that Nhlh1 and Nhlh2 regulate the laterality of commissural axons by transactivating Robo3 expression. To our knowledge, this is the first study to show that a common gene regulatory program operates upstream of Robo3 to regulate commissural axon projection. Our study has brought to the fore a pair of relatively under-explored bHLH molecules, which should serve as a key to deciphering the mechanisms that control the precise and balanced production of commissural neurons.

### Nhlh1 and Nhlh2, as transcriptional activators of Robo3 expression

We showed that Nhlh1 and Nhlh2 function as transcriptional activators in Robo3 induction and that the R6 domain is crucial for their function. Nhlh1/2 belong to the TAL1/SCL subfamily of class II bHLH TFs (Begley et al., 1992; Brown et al., 1992). As with all bHLH TFs, the HLH domain mediates protein interactions, and the basic domain directs DNA binding (Bertrand et al., 2002; Dennis et al., 2019). However, Nhlh1/2 has a unique basic domain containing a highly charged arginine-rich R6 domain that immediately precedes the canonical basic domain. The expanded basic domain of Nhlh1/2 was speculated to enable them to bind to a unique set of regulatory elements (Begley et al., 1992; Brown and Baer, 1994; Brown et al., 1992), although the importance of the R6 domain has not been tested. Our result regarding the requirement of the R6 domain for Robo3 induction suggests that Nhlh1/2 activates Robo3 by binding to specific regulatory motifs that are unlikely to be regulated by other families of bHLH TFs. The HLH domain of Nhlh1/2 is also indispensable for Robo3 induction, because the *Nhlh1* mutant allele lacking the HLH domain fails to activate Robo3. Previous in vitro studies have shown that mouse and human Nhlh1/2 can interact with a range of cofactors, including class I bHLH proteins (E12, E47), LIM-only proteins (Lmo1-4), signal transducer and activator of transcription 3 (STAT3), or even homodimerize (Aoyama et al., 2005; Bao et al., 2000; Brown and Baer, 1994; Fox and Good, 2008; Isogai et al., 2011; Kruger et al., 2004; Manetopoulos et al., 2003; Uittenbogaard et al., 1999).

Heterodimerization between Nhlh1 and Nhlh2, however, is not required for Robo3 induction in vivo, because *Nhlh1* and *Nhlh2* single mutant shows normal commissure formation. We envisage that Nhlh1- and Nhlh2-containing transcriptional complexes function redundantly, by utilizing the same, or different binding sites on the promoter/enhancer, for Robo3 induction. Nhlh1/2 have been suggested to act either as an activator or repressor (Good and Braun, 2013; Isogai et al., 2011; Kruger et al., 2004; Manetopoulos et al., 2003; Wankhade and Good, 2011), perhaps depending on the nature of the cofactors they associate with. Here, we have shown using VP16- and EnR-fusions that Nhlh1/2 serve as activators with respect to Robo3 induction.

### Nhlh1 and Nhlh2, determine the axon projection laterality of commissural neurons

Nhlh1 and Nhlh2 are expressed in post-mitotic immature neurons (Begley et al., 1992; Gobel et al., 1992; Murdoch et al., 1999), therefore, they are expected to regulate early neuronal differentiation. However, their function in neural development remained largely obscure, perhaps in part due to their highly overlapping expression patterns and functional redundancy. In the single mutant, the only defect found during CNS development is the migration of GnRH neurons in *Nhlh2* KO (Kruger et al., 2004).

Double KO of *Nhlh1/2* has been generated and analyzed (Kruger and Braun, 2002; Kruger et al., 2004; Schmid et al., 2007). Surprisingly, although these mice die at birth (Cogliati et al., 2002; Kruger and Braun, 2002), only two defects have been found thus far: an exacerbated GnRH migration defect, and a migration defect of PN neurons causing lateralized PN positions (Kruger et al., 2004; Schmid et al., 2007). Neither of these defects could account for the perinatal lethality, suggesting that there are defects not yet discovered.

By generating *Nhlh1/2* deficient mice, we found that they lack ventral commissures from the spinal cord to the midbrain. The commissural defect provides a satisfactory explanation for the perinatal lethality of these mice, as de-synchronization of the respiratory oscillator owing to disrupted commissural connections in the hindbrain preBötzinger complex has been suggested as a cause of perinatal death in *Robo3* KO mice (Bouvier et al., 2010; Sabatier et al., 2004). The PN migration defect, can also be explained by a lack of attraction of PN leading processes to the FP due to a marked reduction of Robo3 in PN neurons, thereby, failing to pull the PN neuronal soma close to the midline. Besides the obvious lack of ventral commissures, other major axon tracts such as the motor axons, appear normal at the gross level (Figure 6C, data not shown), although a fine-grained analysis of non-commissural axon projections still awaits future investigations as the expression patterns of *Nhlh1/2* extend beyond that of *Robo3*.

We believe that in the commissural system, Nhlh1/2 might be dedicated to controlling Robo3 expression and the absence of commissures in the double mutant is chiefly due to the down-regulation of Robo3. The commissure-less phenotype in the *Nhlh1/2* double mutant and *Robo3* KO mice are remarkably similar (Marillat et al., 2004; Sabatier et al., 2004; Tamada et al., 2008). For example, spinal commissural axons appeared normal in the dorsal half but deflected from the midline upon entering the ventral half of the spinal cord (Figure 6C) (Sabatier et al., 2004). Furthermore, we have provided evidence that the generation and initial axon extension of commissural neurons are not affected in the double mutant.

Are Nhlh1/2 sufficient to confer contralateral axon projection? In the midbrain, most neurons accessible to EP labeling at E11.5 are ipsilaterally-projecting neurons (Inamata and Shirasaki, 2014). We found that some of them could be driven to project contralaterally upon forced expression of Nhlh1/2. This finding is supported by similar observations in the hindbrain. However, the sufficiency of Nhlh1/2 in instructing axon laterality should be considered with caution. Zelina et al. (2014) showed that mouse Robo3, via binding to DCC, promote axon extension to FP by potentiating Netrin-1 responsiveness of commissural axons, which implies that both Robo3 and DCC need to be present for neurons to respond to Netrin-1. We have found no evidence that Nhlh1/2 could induce DCC expression (Zhu, unpublished data), which is reflected by the un-altered DCC levels in the double mutant. It is reasonable to assume that the ectopic contralateral projection driven by Nhlh1/2 force-expression might come from

DCC-expressing ipsilateral neurons. Indeed, DCC IHC on midbrain samples showed that DCC was expressed in the midbrains at E14.5 (Zhu, unpublished data). It might be fair to say that Nhlh1/2 are sufficient to induce contralateral axon projections depending on cellular contexts.

### A global regulatory mechanism for all FP-crossing commissural axons

Whether the specification of shared traits in divergent classes of neurons is determined by common regulatory mechanism or whether they are separately regulated is a fundamental yet unresolved neurodevelopmental issue. The same neurotransmitter phenotype appears to be regulated by different gene regulatory programs that act on distinct cis-regulatory elements in different neuronal types (Hobert and Kratsios, 2019; Lai et al., 2016; Serrano-Saiz et al., 2013). Contralateral axon projection is a common trait imposed during a transient period of development on commissural neurons that are highly heterogeneous in terms of their developmental origin, neurotransmitter types, synaptic targets, and functionality (Chedotal, 2014; Tulloch et al., 2019). Take the mouse spinal cord for example, 8 out of 13 cardinal spinal classes (dI1, dI3, dI4, dI5, dI6, dILA, dILB, and V0), contain a mixture of contralateral and ipsilateral projecting neurons (Alaynick et al., 2011; Lai et al., 2016; Tulloch et al., 2019). Prior to this study, no common regulatory mechanism was found to control Robo3 expression and commissural axon projection (Chedotal, 2014; Friocourt and Chedotal, 2017). A few TFs have been implicated in Robo3 induction, but they function only in specific neuron classes. Lhx2, the only TF confirmed to be an immediate regulator, is responsible for Robo3 expression specifically in dI1 spinal neurons (Ding et al., 2012; Wilson et al., 2008). Accordingly, we found that forced expression of Lhx2 did not induce Robo3 in the rhombic lip and midbrain, suggesting that the key cofactors required for Lhx2 to induce Robo3 are missing. The homeobox domain TF, Dbx1, expressed in the neural progenitor zones, was found to regulate Robo3 expression in the midbrain, and the interneurons in the hindbrain preBötzinger complex (Bouvier et al., 2010; Inamata and Shirasaki, 2014), most likely as an early cell fate determinant. Dbx1 may operate via its downstream target Evx2 to control Robo3 expression in the midbrain (Inamata and Shirasaki, 2014).

Our study found that Nhlh1/2 comprise a global regulatory mechanism that controls contralateral axon projections (Robo3 expression) common to all FP-crossing commissural neurons. Robo3 protein expression levels in the double mutant were barely detectable from the spinal cord to the midbrain, suggesting that transcription of *Robo3* across heterogeneous commissural neuron classes is predominantly driven by an Nhlh1/2-mediated mechanism. Therefore, TFs previously found to function in specific neuron classes are likely to operate via Nhlh1/2. Lhx2, for example, might form a complex with Nhlh1/2, directly or via intermediary cofactors, as a result of which promotes Robo3 transcription. Indeed, in silico analysis has shown that there is an Lhx2 binding site adjacent to the Nhlh1/2 binding site in the highly conserved promoter region (Zhu, unpublished). Dbx1, as a distant upstream regulator, on the other hand, may induce Robo3 expression by activating the expression of Nhlh1/2. We tested this hypothesis by electroporating *Dbx1* into the midbrain and found that concomitant with the induction of Robo3, as previously shown (Inamata and Shirasaki, 2014), *Nhlh2* was indeed induced (Figure S7), supporting this idea.

A recent study showed that the tightly regulated spatial and temporal patterns of Robo3 distribution are highly conserved in the CNS across divergent amniote species (Friocourt et al., 2019). They suggest that deployment of Robo3 to promote contralateral axon projections across the FP might be a strategy that emerged during early vertebrate evolution. The conservation of Robo3 in amniotes raises the possibility that Robo3 may be under the control of conserved gene regulatory programs. Indeed, a phylogenetic analysis showed that *Nhlh1/2* emerged in vertebrates and both genes are present across amniotes (Zhu, unpublished), suggesting that the regulation of Robo3 expression by Nhlh1/2 might be conserved, and has been preserved and incorporated into neuron class specific regulatory programs as neurons diversified during the vertebrate evolution.

### Specificity of commissural neuron fate determination

There is a gross-level spatial and temporal correlation between the expression patterns of *Nhlh1/2* and *Robo3* from the spinal cord to the midbrain. Furthermore, a recent RNA-seq study showed that *Nhlh2* was enriched in Robo3-positive spinal neurons (Tulloch et al., 2019). However, Nhlh1/2 are unlikely to be the sole determinant of commissural fate because their expression pattern appears wider than that of Robo3. For example, in the spinal cord, *Nhlh1/2* are expressed, albeit at lower levels, in what appears to be V1 and V2 spinal neurons which are *Robo3*-negative ipsilateral neurons (Alaynick et al., 2011; Tulloch et al., 2019), and at an earlier stage in young motor neurons (Murdoch et al., 1999). We propose that the specificity of Robo3 expression may be determined by a combination of two levels of regulation. The first is the total amount of Nhlh1 and Nhlh2. The second is an additional layer of Robo3 regulation, via activators or repressors, most likely converging on Nhlh1/2. As discussed above, a range of molecules have been found to be able to interact with Nhlh1/2 in vitro, such as class I bHLH E proteins (E12, E47) and LIM-only proteins (Lmo1-4) (Aoyama et al., 2005; Bao et al., 2000; Brown and Baer, 1994; Isogai et al., 2011; Kruger et al., 2004; Manetopoulos et al., 2003; Uittenbogaard et al., 1999). There are precedents showing that E proteins and Lmo proteins are capable of fine tuning the gene regulatory mechanisms mediated by class II bHLH proteins (Joshi et al., 2009; Le Dreau et al., 2018). It would be of future interest to investigate whether any of these potential cofactors modulate the activity of Nhlh1/2 in the specification of commissural neuronal fate. The intersection of the global mechanism uncovered here with neuron subtype-specific regulatory mechanisms should ultimately determine the specific and balanced production of contralateral- versus ipsilateral-projection neurons to achieve optimized outcomes at the neural circuitry level.

## Supporting information

Supplemental figure all

## ACKNOWLEDGEMENTS

The authors thank Ms. Noriko Yamatani, Ms. Mami Yokoyama, and Ms. Sayuri Noguchi for their technical assistance, Dr. Alexandra Pierani for *Dbx1* expression construct, Dr. Yasuto Tanabe for *nls-EGFP* expression construct, Dr. Masahiko Hibi for *VP16* and *EnR* containing plasmids, Drs. Toshihiko Shiroishi and Tomoko Sagai for *Shh* riboprobe, Drs. Fujio Murakami and Yash Hiromi for critical reading of the manuscript. We would like to thank Editage (www.editage.com) for English language editing. This study was funded by Grants-in-Aid for Scientific Research from the Ministry of Education, Culture, Science and Technology, Japan, contract grant nos: 16K07010, 20K06865 (to YZ), 20H03345 (to TH).

## AUTHOR CONTRIBUTIONS

YZ conceived the study. AM and YZ designed and conducted the experiments, analyzed data and wrote the manuscript. RA and YS generated the mutant mice. TH contributed materials and advices. YZ and TH provided funding. All authors read and approved the manuscript.

## DECLARATION OF INTERESTS

The authors declare no competing interests.

## MATERIALS and METHODS

### Key Resource Table

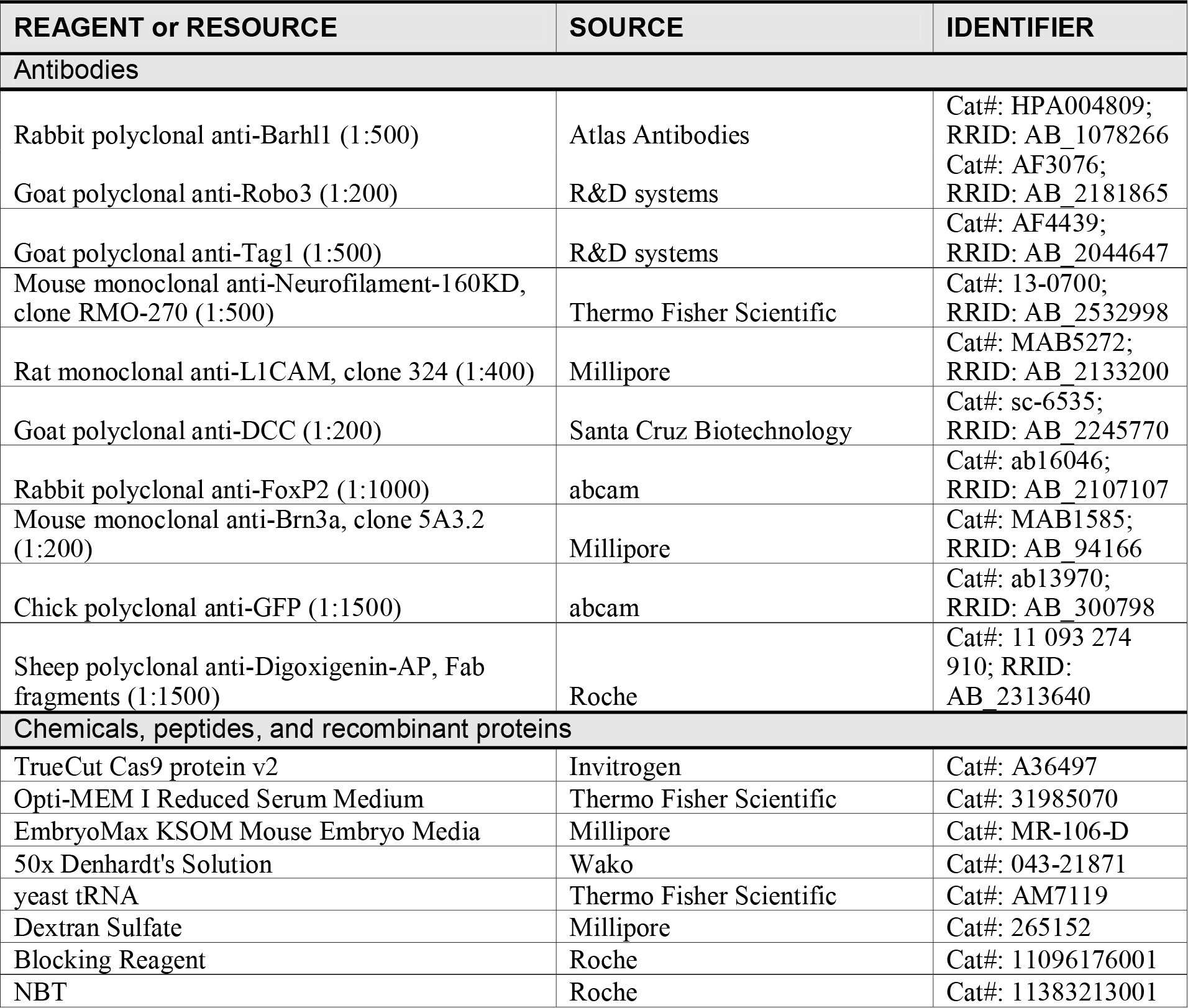
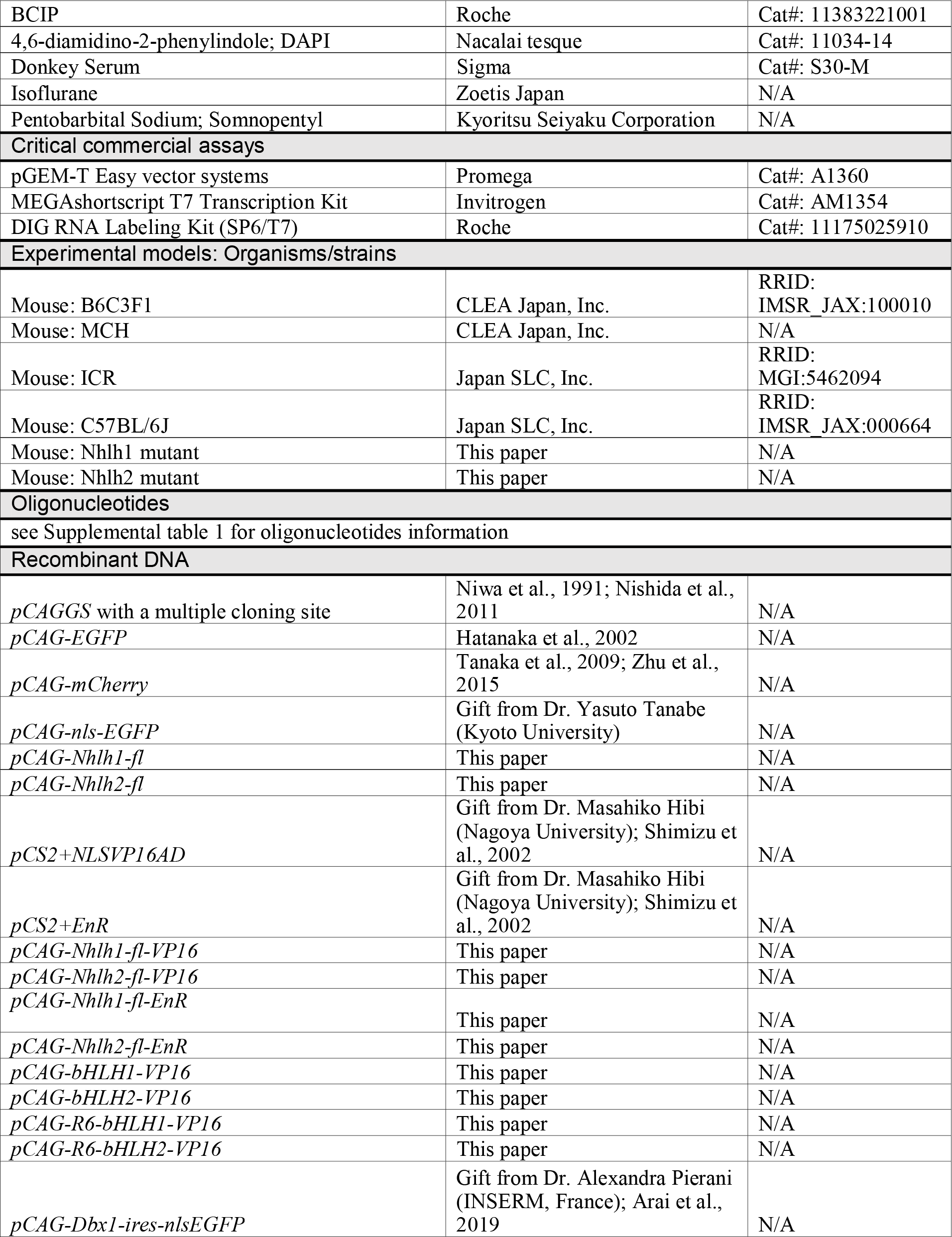
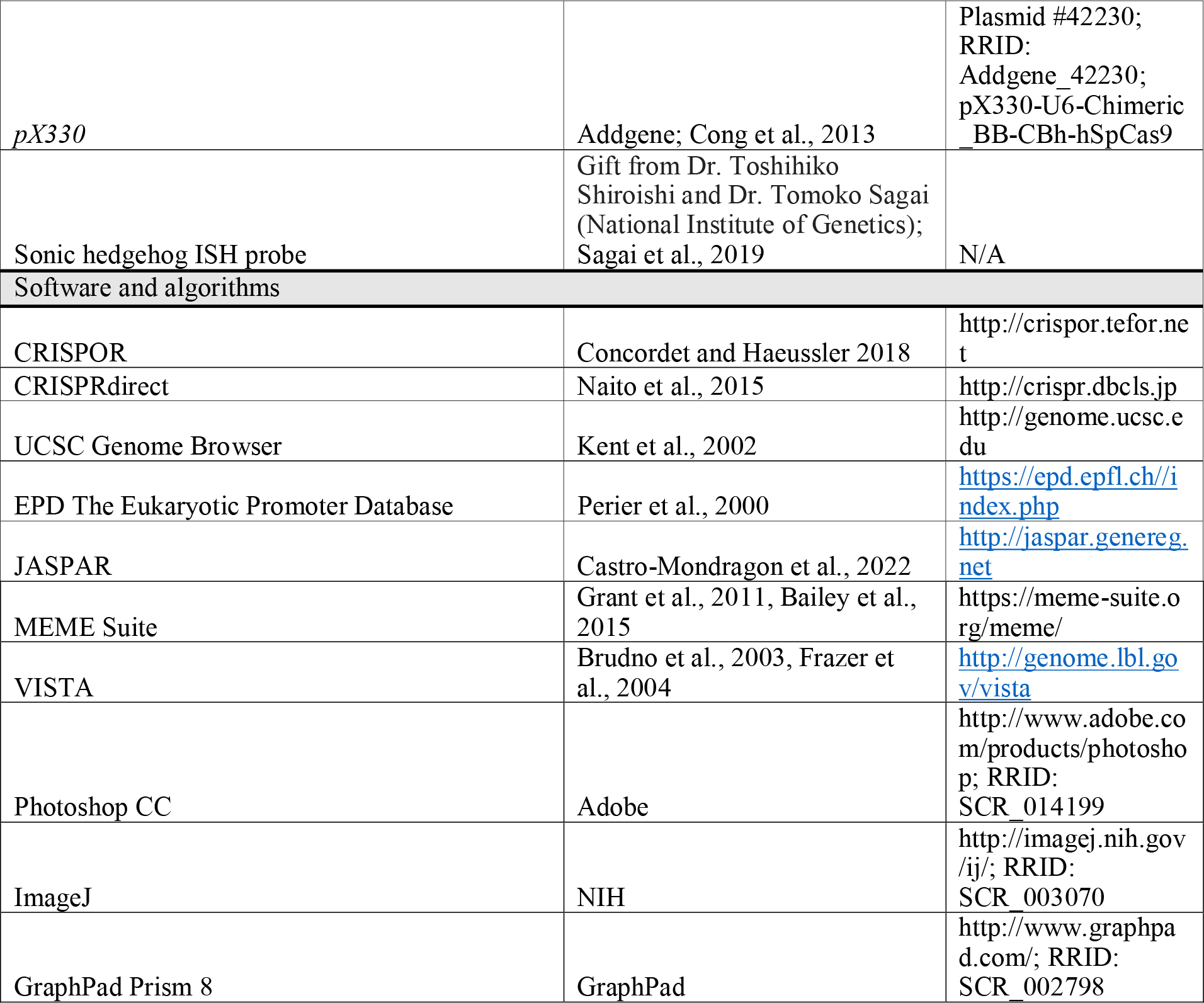

### DNA constructs

Expression constructs *pCAG-EGFP* and *pCAG-mCherry* have been described before (Hatanaka and Murakami, 2002; Tanaka et al., 2009; Zhu et al., 2015).

*pCAG-nls-EGFP* was a kind gift from Dr. Yasuto Tanabe (Kyoto University, Japan). Full-length (fl) *Nnlh1* and *Nhlh2* expression constructs were constructed by PCR amplification of the coding sequences of *Nhlh1* (GenBank accession: NM_010916) and *Nhlh2* (GenBank accession: NM_178777) from mouse E11.5 brain cDNA. The PCR products were cloned into a *pCAGGS* vector with a multiple cloning site inserted (Nishida et al., 2011; Niwa et al., 1991) to generate *pCAG-Nhlh1-fl* and *pCAG-Nhlh2-fl*. The PCR products were also cloned into a *pCAG-2HA* plasmid which contains the *pCAGGS* backbone with two Hemagglutinin (HA) tags. This resulted in the generation of fusion proteins of Nhlh1 and Nhlh2 with two HA tags at their N-termini. The HA fused or non-fused versions of Nhlh1 and Nhlh2 were confirmed to be equivalent in inducing Robo3 expression (data not shown). To produce Nhlh1- and Nhlh2-VP16 fusion proteins, the *VP16* and *EnR* coding sequences were excised from *pCS2+NLSVP16AD* and *pCS2+EnR* plasmids (kind gifts from Dr. Masahiko Hibi, Nagoya University, Japan) (Shimizu et al., 2002), respectively, and cloned into *pCAG-Nhlh1-fl* and *pCAG-Nhlh2-fl* at 5’ to and in frame with *Nhlh1* and *Nhlh2* coding sequences. To construct the *pCAG-bHLH1-VP16* and *pCAG-bHLH2-VP16* constructs, the bHLH domains (excluding the R6) of *Nhlh1* and *Nhlh2* were PCR amplified with forward primer, 5’ATCTCGAGGCCACGGCCAAGT3’, for both *Nhlh1* and *Nhlh2*, and reverse primers 5’ATAGCGGCCGCTCAGACGTCCAGCA3’ for *Nhlh1* and 5’ATAGCGGCCGCCTACACGTCCAGGA3’ for *Nhlh2*. To construct *pCAG-R6-bHLH1-VP16* and *pCAG-R6-bHLH2-VP16*, the R6-bHLH domains, from *Nhlh1* and *Nhlh2*, were PCR amplified with forward primers 5’CGCTCGAGCACTTGAGTCGTGAG3’ and 5’ATCTCGAGCAGCTGAGCCGCGAA3’, for *Nhlh1* and *Nhlh2*, respectively, and the same reverse primers as above. The PCR fragments were cut with XhoI and NotI, and were subcloned into XhoI and NotI digested *pCAG-Nhlh1-VP16* and *pCAG-Nhlh2-VP16* plasmids to replace the full-length Nhlh1 and Nhlh2, respectively. The *Dbx1* expression construct (*pCAG-Dbx1-ires-nlsEGFP*) was a kind gift from Dr. Alexandra Pierani (INSERM, France) (Arai et al., 2019). All the cloned expression constructs were confirmed by sequencing.

Constructs for generating riboprobes to detect *Robo3*, *Nhlh1*, *Nhlh2*, *Netrin-1* were made using the pGEM-T Easy vector system (Promega) by PCR amplification from E11.5 mouse brain cDNA with the following primer pairs: 5’ACAGCAGCCTATCTAGGCCA3’ and 5’TCTGGTATTCAGTGATGACCCC3’ for *Robo3*, 5’TGTTCAGCCACAAGCTGC3’ and 5’GCGCTCCTCACGACTCAA3’ for *Nhlh1*, 5’CTGCCAAAGGCGACTCAT3’ and 5’AGACGGGGTGTTTTTGGA3’ for *Nhlh2*, and 5’CTTCCTCACCGACCTCAATAAC3’ and 5’TAGAGCTCCATGTTGAATCTGC3’ for *Netrin-1*. Sonic hedgehog (*Shh*) ISH probe was a kind gift from Dr. Toshihiko Shiroishi and Dr. Tomoko Sagai (National Institute of Genetics, Japan) (Sagai et al., 2019).

For generating sgRNA for gene editing in fertilized eggs, oligonucleotide pairs were annealed and cloned into BbsI digested *pX330* (Addgene, Plasmid#42230) (Cong et al., 2013): for *Nhlh1-sg1*, 5’CACC<u>GGACCGGGCCCCGATGGTGC</u>3’ and 5’AAAC<u>GCACCATCGGGGCCCGGTCC</u>3’; for *Nhlh1-sg2*, 5’CACC<u>GCTAGGTTGAAGGCTTCCACG</u>3’ and 5’AAAC<u>CGTGGAAGCCTTCAACCTAGC</u>3’; for *Nhlh2-sg1*, 5’CACC<u>GCGAAGAAAAGCGCCGCCGC</u>3’ and 5’AAAC<u>GCGGCGGCGCTTTTCTTCGC</u>3’; for *Nhlh2-sgRNA2*, 5’CACCG<u>TTCTTGTCCGGAGGCAGCGT</u>3’ and 5’AAAC<u>ACGCTGCCTCCGGACAAGAA</u>C3’. The target regions are underlined.

### Generation of *Nhlh1* and *Nhlh2* mutant mice

The CRISPR/Cas9 guide sequences for *Nhlh1* and *Nhlh2* genes were selected using CRISPOR (http://crispor.tefor.net) (Concordet and Haeussler, 2018) and CRISPRdirect (http://crispr.dbcls.jp) (Naito et al., 2015) webtools (Figure S3A). The sgRNAs were synthesized as reported previously (Ajima et al., 2017). B6C3F1 (C57BL/6N X C3H/HeN) female mice were super-ovulated and mated with B6C3F1 males, and fertilized eggs were collected from oviducts. The 50 ng/μl synthesized sgRNAs and 100 ng/μl TrueCut Cas9 protein v2 (Invitrogen) were premixed in Opti-MEM (Thermo Fisher Scientific) and electroporated into fertilized eggs using CUY21EDIT II electroporator and LF501PT1-10 platinum plate electrode (BEX Co. Ltd.) following the method reported previously (Hashimoto and Takemoto, 2015). The electroporated zygotes were cultured in KSOM (Millipore) at 37L under 5% CO_2_ until the two-cell stage after 1.5 days. Thereafter, 20 to 32 two-cell stage embryos were transferred into the uterus of pseudo-pregnant MCH females at 0.5 days post coitum.

Founder mice were screened for edited Nhlh1 and Nhlh2 alleles which were subsequently sequenced to obtain the precise nature of the edited alleles. Two founder lines were then chosen for breeding, backcrossing and analyses (Figure S3C, D).

### Animals

For expression studies with ISH and in utero EP, timed pregnant ICR mice (Nihon SLC, Shizuoka, Japan) were used. Noon of the day on which a vaginal plug was detected was designated as embryonic day (E) 0.5. Nhlh1 and Nhlh2 mutant mouse lines, initially on a B6C3F1 hybrid background, were backcrossed to C57BL/6J for at least two to five generations before phenotype analyses. Double and single heterozygotic colonies were maintained. To generate double homozygotic embryos, double heterozygotic male and female mice were crossed. Genotyping were performed on tail lysis using the following primers. For *Nhlh1* mutant, 5’AGTCGGGCTTTAGCGACTGT3’ and 5’GCGAAGGCTAGGTTGAAGG3’ were used; and for *Nhlh2* mutant 5’AAACTACTACCCACGCTGCC3’, 5’CCACTACTCCTTGCAAATCAAGA3’, 5’GACTAGAAAGAACAGGCTGCAAA3’, and 5’AGACGTTCTTGTCCGGAGTAG3’ were used. All animal maintenance and manipulations were carried out in accordance with the Guidelines for the Welfare and Use of Laboratory Animals of the National Institute of Genetics, Japan.

### In utero electroporation

In utero EP was performed essentially as previously described (Zhu et al., 2020) with some modifications. The pregnant mice were anesthetized with a combination of isoflurane (1.0% in air) and Pentobarbital Sodium (Somnopentyl, Kyoritsu Seiyaku Corporation, Tokyo, Japan, 80 mg/kg body weight). The uterus was exposed after abdominal incision and approximately 2 μl of plasmid was injected into the IV ventricle or the cerebral aqueduct of E12.5 or E11.5 embryos. Five square electric pulses (30V, 50 ms duration at 200 ms intervals) were applied using a forceps-type electrode (CUY650P5 or CUY650P2, Nepa Gene, Japan) connected to a square-pulse generator (CUY21, BEX, Japan).

### Tissue processing and sectioning

Pregnant mice were killed by cervical dislocation and embryos were taken out from the uterus. Mouse brainstems were dissected out in phosphate-buffered saline (PBS, pH7.4) and fixed in 4% paraformaldehyde (PFA, 0.01 M PBS) at 4°C for 6-7 hr. The tissues were then cryo-protected in 30% sucrose (in PBS) overnight at 4°C and embedded in O.C.T compound (Sakura FineTek, Japan). Frozen sections of 20 μm thickness were obtained with a cryostat (Leica CM3050S).

### In situ hybridization on sections

Tissue sections were pre-hybridized at 65°C for 2 hr in hybridization buffer comprised of: 50% Formamide (deionized and nuclease tested, Nacalai tesque), 1.3x SSC, 1xDenhardt’s Solution (Wako), 0.1mg/ml yeast tRNA and 10% (w/v) Dextran Sulphate (M.W.500000, Millipore). The sections were then incubated with hybridization buffer containing 1.5μg/ml of anti-sense riboprobe at 65°C overnight. The next day, sections were washed twice at 65°C with formamide wash buffer (50% Formamide, 1xSSC, 0.1% Tween-20), and then twice at room temperature (RT) with 1xMABT (100mM Maleic Acid, 150mM NaCl, 0.1% Tween-20). Blocking was performed in 1xMABT with 2% Blocking Reagents (Roche) and 10% donkey serum (Sigma) for 1 hr at RT, and reacted with anti-DIG-AP Fab fragments (1:1500, Roche) in the blocking solution at 4°C overnight. The next day, slides were washed three times in 1xMABT and rinsed once in NTMT solution (0.1M Tris-HCl pH9.5, 0.1M NaCl, 0.05M MgCl2, 0.1% Tween-20), and then color developing was carried out in NTMT containing NBT (100ug/ml) and BCIP (50ug/ml).

### Immunohistochemistry

IHC was performed as previously described (Zhu et al., 2015). The sections were blocked with 10% donkey serum (Sigma) in PBSTx (0.2% Triton X-100) for 1 hr at RT followed by incubation with primary antibodies at 4°C overnight. After washing with PBSTx, the sections were then incubated with the secondary antibodies at RT for 2 hr. Slides were counter-stained with 0.03% 4,6-diamidino-2-phenylindole (DAPI, Nacalai tesque). The primary antibodies used were: rabbit anti-Barhl1 polyclonal antibody (Atlas Antibodies, HPA004809, Sigma, 1:500), goat anti-Robo3 polyclonal antibody (R&D Systems, AF3076, 1:200), goat anti-Tag1 polyclonal antibody (R&D Systems, AF4439, 1:500), mouse anti-Neurofilament-160KD monoclonal antibody (clone: RMO-270, Zymed, 13-0700, 1:500), rat anti-L1CAM monoclonal antibody (clone 324, Chemicon, MAB5272, 1:400), goat anti-DCC polyclonal antibody (Santa Cruz Biotechnology, sc-6535, 1:200), rabbit anti-FoxP2 polyclonal antibody (abcam, ab16046, 1:1000), mouse anti-Brn3a monoclonal antibody (clone: 5A3.2, Millipore, MAB1585, 1;200), chick anti-GFP polyclonal antibody (abcam, ab13970, 1:1500). The secondary antibodies used were Cy3-donkey anti-rabbit IgG (Jackson ImmunoResearch, 1:300) for Barhl1, FoxP2 antibodies, Cy3-donkey anti-goat IgG (Jackson ImmunoResearch, 1:300) for Robo3, Tag1, DCC antibodies, Cy3-donkey anti-mouse IgG (Jackson ImmunoResearch, 1:300) for Neurofilament, Brn3a antibodies, Cy3-donkey anti-rat IgG (Jackson ImmunoResearch, 1:300) for an L1CAM antibody, Alexa Fluor 488-donkey anti-chick IgY (Jackson ImmunoResearch, 1:400) for a GFP antibody.

### Transcriptional Factor binding site analysis

Promoter sequences of mouse, human, rabbit, cow, ferret and opossum *Robo3* were extracted using the UCSC genome browser (http://genome.ucsc.edu) (Kent et al., 2002). The transcriptional start site (TSS) information were obtained from the Eukaryotic Promoter Database (EPD, https://epd.epfl.ch//index.php) (Perier et al., 2000). The Nhlh1 and Nhlh2 binding site position frequency matrices (PFM) were downloaded from JASPAR (http://jaspar.genereg.net) (Castro-Mondragon et al., 2022). Using the FIMO tool from the MEME suite (https://meme-suite.org/meme/, (Bailey et al., 2015; Grant et al., 2011), the promoter region of mouse *Robo3* was scanned for Nhlh1 and Nhlh2 binding sites with their respective PFM from JASPAR. Only regions identified to be bound by both Nhlh1 and Nhlh2 were illustrated in Figure 1. Phylogenetic foot printing was performed by using the mVISTA tool in VISTA (http://genome.lbl.gov/vista) (Brudno et al., 2003; Frazer et al., 2004). Conservation parameters were set to be: minY 50%, minID 70%, min Length 100 bases.

### Image acquisition and processing

Fluorescence images on sections or flat-mounted brainstems were captured with a CCD camera (Axiocam, Zeiss) attached to an upright microscope (BX-60, Olympus) at 1296x1030 pixel resolution. Objective lens used were: 2x Plan Apo with numerical aperture (NA) 0.08 (Olympus), 4x UPlan Apo with NA 0.16 (Olympus), 10x UPlan Apo with NA 0.40 (Olympus) and 20x UPlan Apo with NA 0.70 (Olympus). All bright field images on ISH sections, as well as some fluorescence images were captured with an All-in-one Fluorescence Microscope (BZ-X700, Keyence) with an objective lens of CFI 10x Plan Apo Lamda, NA 0.45 (Nikon). Adobe Photoshop CC was used to adjust contrast and brightness of images and to assemble figures.

### Quantification and statistical analysis

For quantifying the commissural index in the electroporated midbrains, fluorescence images of the whole-mounted brainstems were imported into ImageJ (NIH, http://imagej.nih.gov/ij/). An ROI of defined size over the ipsilateral extending axons, the contralateral extending axons and a background region on the sample were measured for its mean fluorescence level denoted as F_i_, F_c_, and F_b_, respectively. The commissural index was calculated as (F_c_-F_b_)/(F_i_-F_b_). For quantifying the commissural index in the electroporated hindbrains, fluorescence images of the hindbrain sections after IHC were imported into ImageJ. An ROI of defined size over a region just below the rhombic lip, the FP region, and a background region on the sample were measured for its mean fluorescence level denoted as F_ep_, F_fp_, and F_b_, respectively. The commissural index was calculated as (F_fp_-F_b_)/(F_ep_-F_b_). The quantified data were represented by scatter plots with median and upper and lower quartiles indicated and statistical analyses were performed by Mann-Whitney U test using Prism 8 (GraphPad). To quantify the number of Brn3a positive neurons on the spinal cord samples, the number of Brn3a signal positive cells were manually counted on 2-3 sections for each sample and subsequently averaged. This was done for all the samples on spinal cord sections of approximately equivalent axial levels. The data points were represented on a scatter plot using Microsoft excel.

## Notes

### Competing Interest Statement

The authors have declared no competing interest.

### Summary of Updates

The revised version contains 7 instead of 8 figures. Figure 6 and 7 of previous version are now combined into one figure. Some of the supplemental figures in the previous version are combined, hence, the number of supplemental figures are reduced from 16 to 7.

