## Supplemental figure all for "Nhlh1 and Nhlh2, a global transcriptional mechanism regulating commissural axon projection via Robo3 activation"

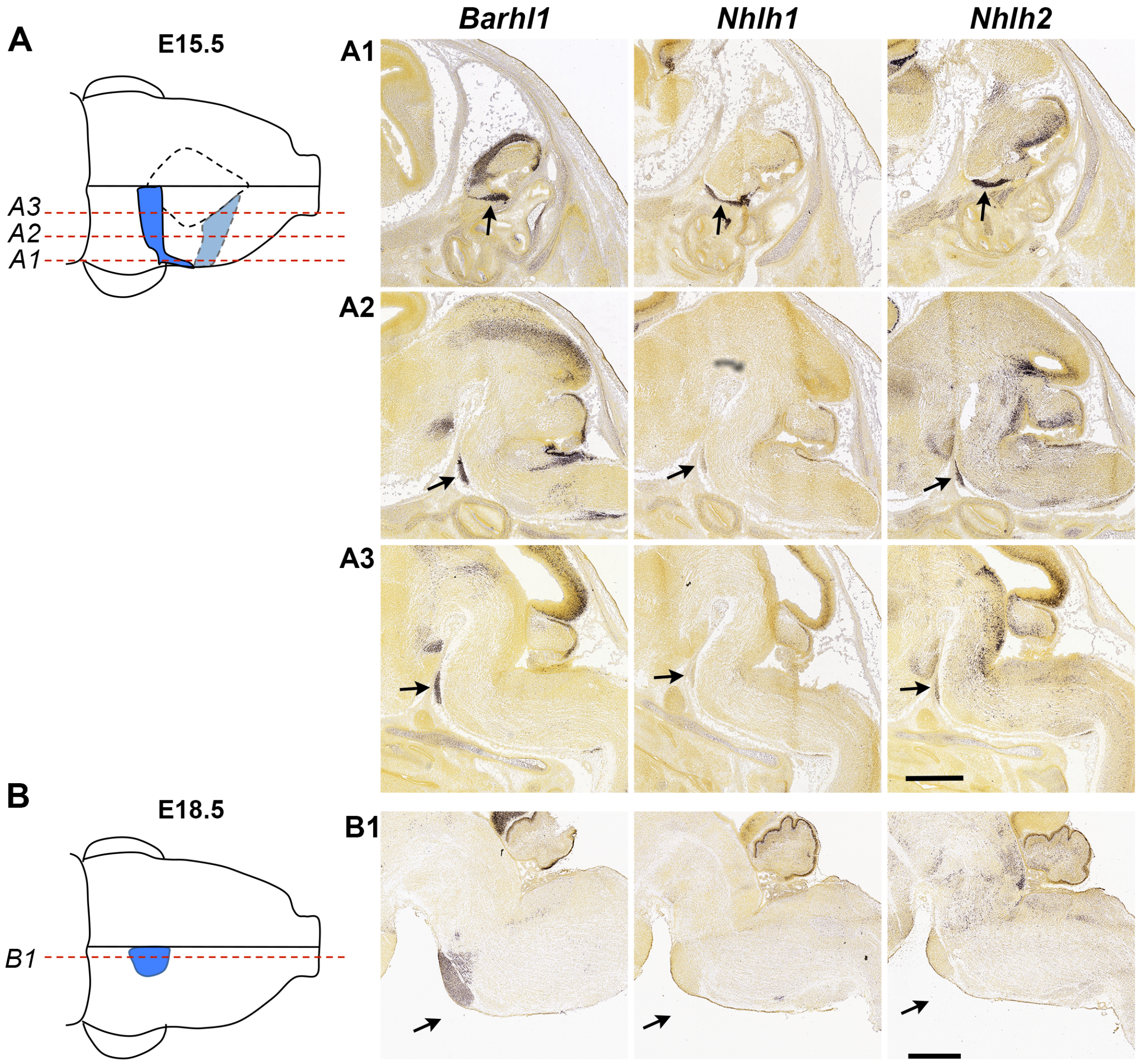


**Figure S1. Expression of Nhlh1 and Nhlh2 in pre-crossing, and their down-regulation in post-crossing, PN neurons.** **(A)** A schematic of a ventral view of an E15.5 hindbrain with the migratory stream of PN neurons indicated in blue. The red dashed lines indicate the approximate mediolateral positions of the three sets of sagittal sections shown in (A1), (A2) and (A3). **(B)** A schematic of a ventral view of an E18.5 hindbrain when most PN neurons have entered the nuclear region and become post-crossing. The red dashed line indicates the approximate position of the set of sagittal sections shown in (B1). **(A1), (A2), (A3) & (B1)** ISH images selected from the Allen Developing Mouse Brain Atlas (https://developingmouse.brain-map.org/). PN neurons are marked by Barhl1 ISH (left panel), and Nhlh1 (mid panel) and Nhlh2 (right panel). ISH images were aligned with that of Barhl1 on sections of similar mediolateral levels. Arrows indicate the PN neurons. Scale bars: 800µm in (A1), (A2) & (A3); 800 µm in (B1). **Related to Figure 1.**

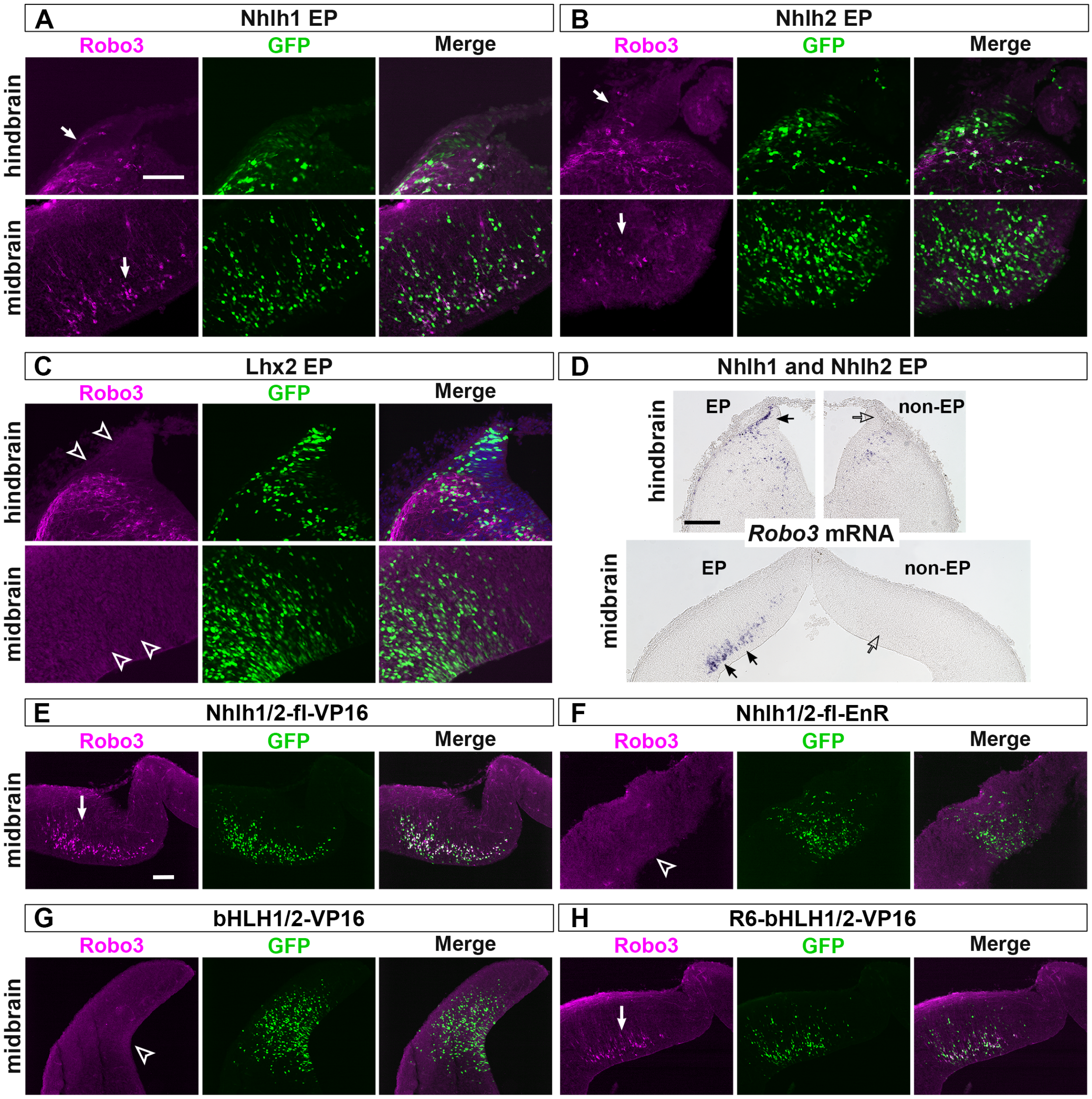


**Figure S2. Testing the forced expression of various constructs in ectopic Robo3 induction.** For all experiments in this figure, the expression construct indicated above each panel was electroporated together with nls-EGFP into the rhombic lip of the hindbrain or into the midbrain at E12.5 and the samples were analyzed for Robo3 induction at E14.5. **(A) & (B)** Full length Nhlh1 or Nhlh2 alone induced ectopic Robo3 expression in the hindbrain (n=3 for Nhlh1, and n=2 for Nhlh2) and the midbrain (n=3 for Nhlh1, and n=2 for Nhlh2), although the induction appeared weaker than Nhlh1 and Nhlh2 combined EP. **(C)** Lhx2 EP did not induce ectopic Robo3 expression (n=3 for the hindbrain, n=2 for the midbrain). **(D)** Nhlh1 and Nhlh2 EP induced Robo3 transcription as shown by Robo3 ISH. **(E), (F), (G) & (H)** EP of VP16- and EnR-fusion constructs in the midbrain. Nhlh1/2-fl-VP16 (n=2) and R6-bHLH1/2-VP16 (n=2) induced ectopic Robo3 expression, while Nhlh1/2-fl-EnR (n=2) and bHLH1/2-VP16 (n=2) did not. Scale bars: 100 µm in (A), (B) and (C); 200 µm in (D); 100 µm in (E), (F), (G) and (H). **Related to Figure 2 and Figure 3.**

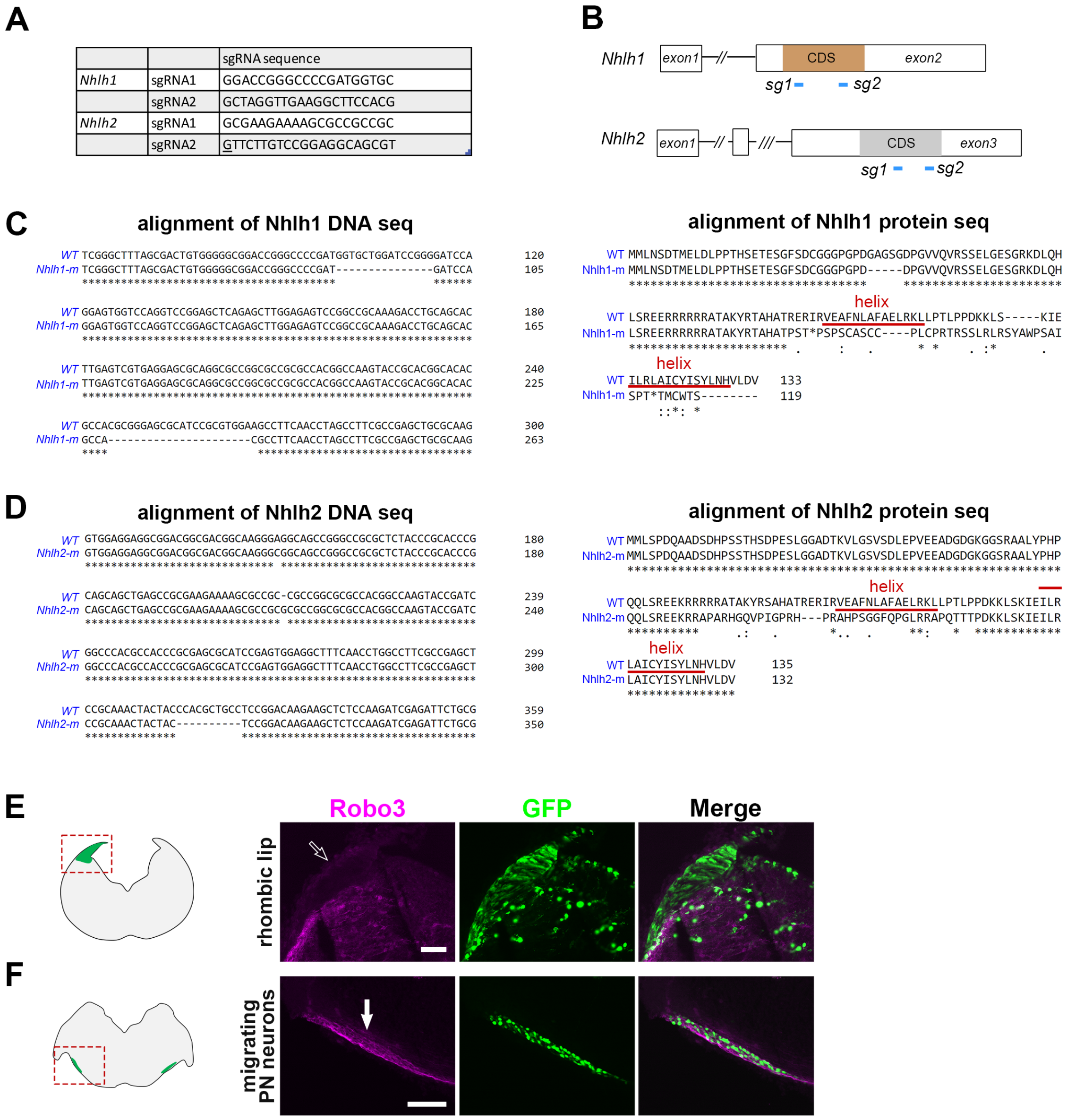


**Figure S3. Generation of Nhlh1 and Nhlh2 mutant alleles by CRISPR-Cas9 gene editing in fertilized eggs.** **(A)** Sequences of the two sgRNAs used for Nhlh1 and Nhlh2 alleles, respectively. **(B)** A schematic showing the approximate positions of the two sgRNAs on the Nhlh1, or Nhlh2 locus. **(C) & (D)** Alignment of DNA sequences and protein sequences between the wild type and Nhlh1 mutant allele, and between the wild type and Nhlh2 mutant allele, respectively. **(E)** EP of constructs expressing Nhlh1 and Nhlh2 mutant proteins in the hindbrain did not induce ectopic Robo3 expression within the rhombic lip (n = 2). **(F)** EP of constructs expressing Nhlh1 and Nhlh2 mutant proteins in the migrating PN neurons did not suppress their Robo3 expression (n = 2). Scale bars: 100 µm in (E); 100 µm in (F). **Related to Figure 6, Figure 7.**

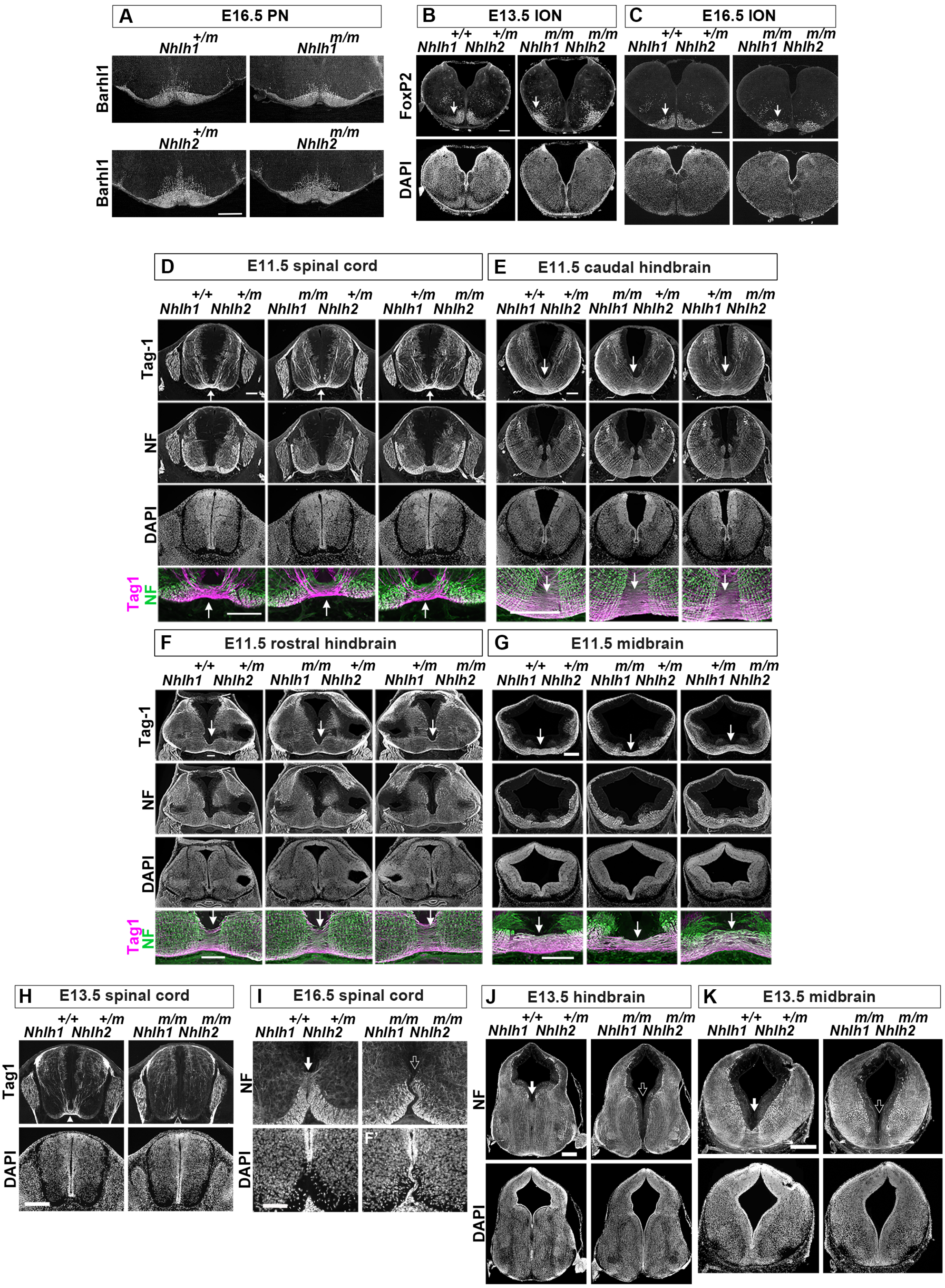
A E16.5 PN
Nhlh1+/m
Nhlh1m/m
Nhlh2+/m
Nhlh2m/m
Barhl1
B E13.5 ION
C E16.5 ION
Nhlh1+/+ Nhlh2+/m
Nhlh1m/m Nhlh2m/m
FoxP2
DAPI
D E11.5 spinal cord
E E11.5 caudal hindbrain
F E11.5 rostral hindbrain
G E11.5 midbrain
Nhlh1+/+ Nhlh2+/m
Nhlh1m/m Nhlh2+/m
Nhlh1+/m Nhlh2m/m
Tag-1
NF
DAPI
Tag1
NF
H E13.5 spinal cord
I E16.5 spinal cord
J E13.5 hindbrain
K E13.5 midbrain
Nhlh1+/+ Nhlh2+/m
Nhlh1m/m Nhlh2m/m
Tag1
NF
DAPI
F'

**Figure S4. Nhlh1 and Nhlh2 mutant phenotype analysis. (A)** PN formation was not affected in Nhlh1 and Nhlh2 single mutant. PN formation was analyzed by Barhl1 IHC on E16.5 rostral hindbrain sections. PN position, shape and size appeared comparable between the heterozygotes and homozygotes of Nhlh1 mutant (n = 3 for each geneotype) and Nhlh2 mutant (n = 2 for each genotype). **(B) & (C)** ION is lateralized in Nhlh1 and Nhlh2 double mutant. IO neurons were labelled by FoxP2 IHC at E13.5 (B) and E16.5 (C). IO neurons were positioned at a distance away from the ventral midline in the double mutant (n = 2 for E13.5, and n = 2 for E16.5), whereas in the control genotype, they were in close juxtaposition to the ventral midline (n = 2 for E13.5, and n = 2 for E16.5). **(D), (E), (F) & (G)** Commissural formation appeared unaffected in Nhlh1 and Nhlh2 single mutant. Commissural axons were labelled by Tag1 IHC and the overall axonal patterns were labelled by NF IHC. Ventral commissures appeared comparable between the control geneotype of $Nhlh1^{+/+}Nhlh2^{+/m}$ (n = 2), and the single mutant $Nhlh1^{m/m}Nhlh2^{+/m}$ (n = 2) and $Nhlh1^{+/m}Nhlh2^{m/m}$ (n = 2), in the spinal cord (D), caudal hindbrain (B), rostral hindbrain (C), and midbrain (D). Higher magnification images of the commissural region across the ventral midline are shown in the bottom panels. **(H), (I), (J) & (K)** Commissure-less phenotype in the Nhlh1 and Nhlh2 double mutant persisted into later developmental stages. Commissure formation was analyzed either by Tag1 or NF IHC. Lack of commissure formation continued to be observed in E13.5 spinal cord (H) (n = 2 for each genotype), E16.5 spinal cord (I) (n = 2 for each genotype), E13.5 hindbrain (J) (n = 2 for each genotype) and E13.5 midbrain (K) (n = 2 for each genotype). PN: pontine nucleus; ION: inferior olive nucleus. Scale bars: 200 μm in (A); 200 μm in (B) and (C); 100 μm in (D); 200 μm in (E), (F) and (G); 200 μm in (H); 100 μm in (I); 400 μm in (J) and (K). **Related to Figure 6.**

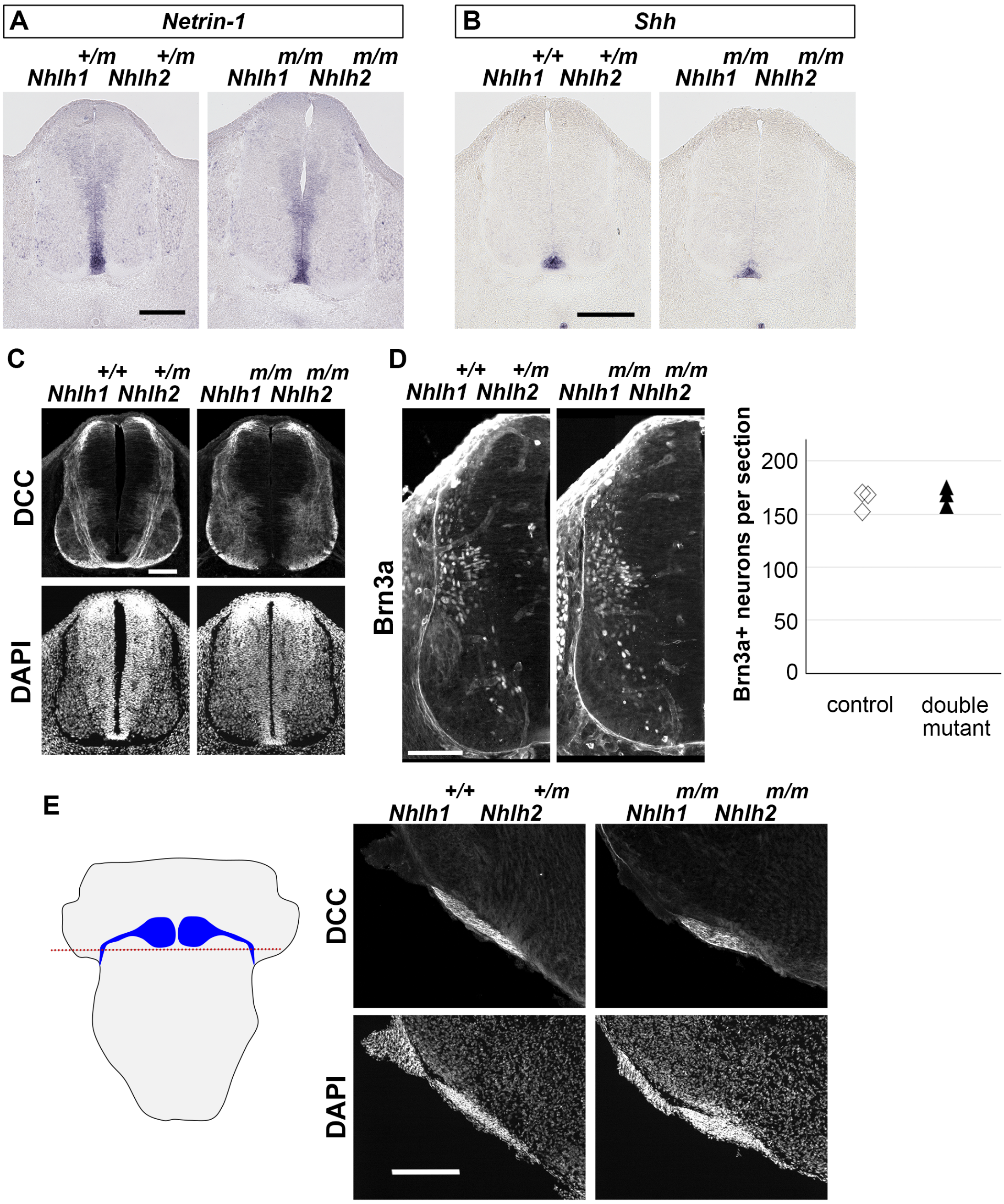


**Figure S5. Expression of Netrin-1, Shh, DCC and Brn3a were not affected in Nhlh1 and Nhlh2 double mutant.** **(A) & (B)** Expression of Netrin-1 and Shh were examined by ISH on E11.5 spinal cord sections, and appeared comparable between the control and the dboule mutant. **(C)** DCC expression pattern was similar between the control and the double mutant in the dorsal half of the spinal cord where most spinal commissural neurons reside (n = 3 for each geneotype). **(D)** Spinal cord sections of control and double mutant were subjected to Brn3a IHC. Brn3a expressing spinal neurons which encompassing several classes of both ipsi-lateral and commissural dorsal spinal neurons were similar in numbers between the control (n = 3) and the double mutant (n = 3) as shown by the quantification depicted in the scatter plot. **(E)** DCC IHC on an E16.5 hindbrain section from approximately the level indicated in the schematic on the left. DCC was expressed at a smilar level between the control (n = 2) and the double mutant (n = 2). Scale bars: 200 μm in (A); 200 μm in (B); 100 μm in (C); 100 μm in (D); 200 μm in (E). **Related to Figure 6**.

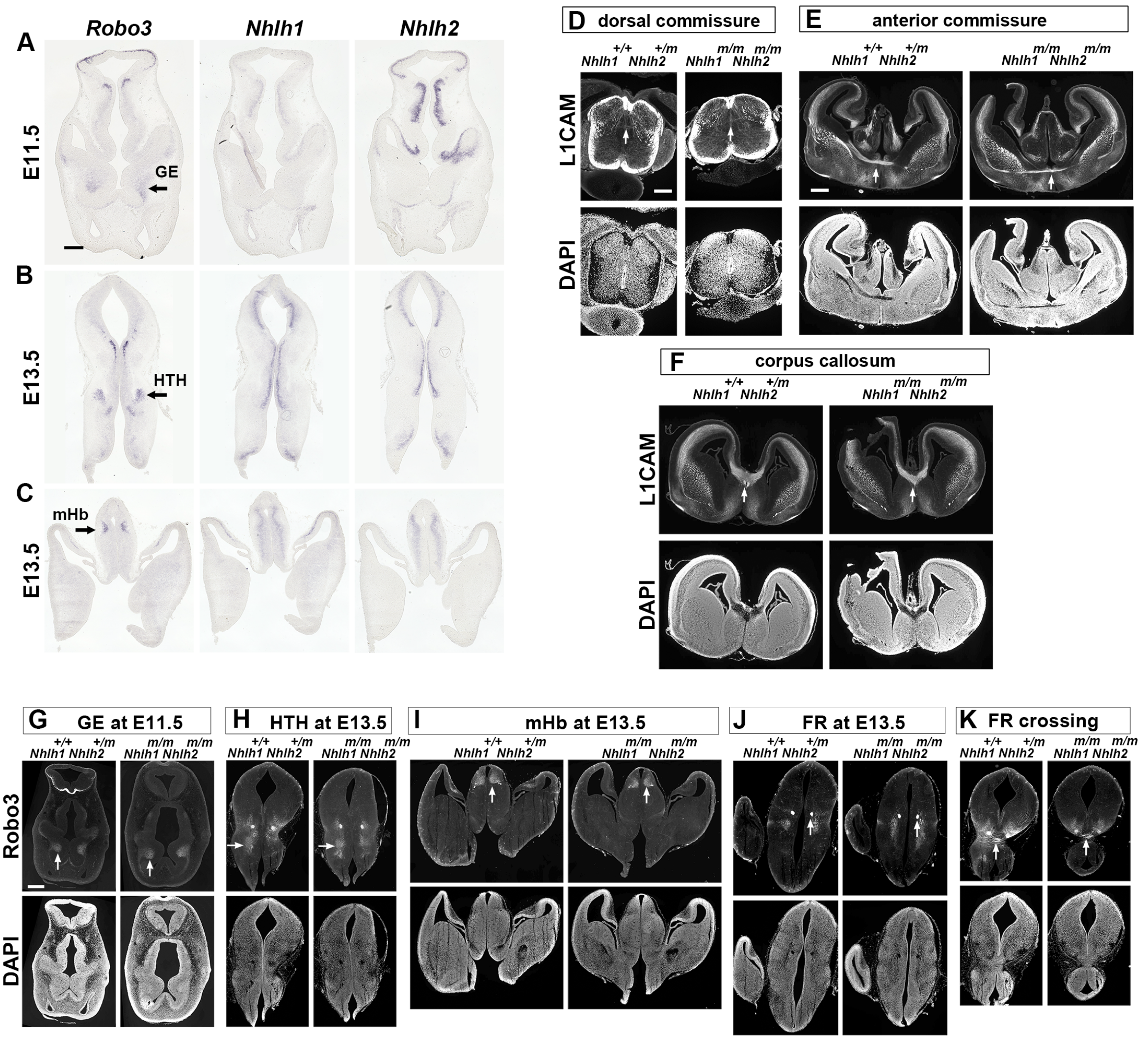


**Figure S6. Expression pattern, phenotype analyses, and Robo3 down-regulation analysis in the forebrain or non-FP crossing commissures. (A), (B) & (C)** Robo3 expression is not well correlated with those of Nhlh1 and Nhlh2 in brain structures rostral to the midbrian. **(A)** Robo3 is expressed in GE at E11.5, but not Nhlh1 and Nhlh2. **(B)** Robo3-positive neurons in HTH at E13.5 do not appear to express Nhlh1 and Nhlh2. **(C)** mHb neurons in the E13.5 epithalamus express Robo3, but they either do not or only partially express Nhlh1 and Nhlh2. **(D), (E) & (F)** Commissural formation not via FP is unaffected in Nhlh1 and Nhlh2 double mutant. The dorsally crossing commissural axons in the spinal cord (D), the anterior commissure (E), and the corpus callosum (F) in the forebrain appeared comparable between the control and the double mutant (n = 2 for each genotype). **(G), (H), (I), (J) & (K)** Robo3 expression is not affected in brain structures rostral to the midbrain in the Nhlh1 and Nhlh2 double mutant. Robo3 expression was examined by IHC. Robo3 expression level was comparable between the control and the double mutant, in the GE (G) (n = 2 for each genotype), the HTH (H) (n = 2 for each genotype), the mHb (I) (n = 2 for each geneotype), the caudally extending FR (J) (n = 2, for each genotype) and the FR crossing the ventral midline at the midbrain/hindbrain junction (K) (n = 2 for each genotype). GE: ganglionic eminence; HTH: hypothalamus; mHb: medial habenular nucleus; FR: fasciculus retroflexus. Scale bars: 200 μm in (A), (B) and (C); 200 μm in (D); 400 μm in (E) and (F); 400 μm in (G), (H), (I), (J) and (K). **Related to Figure 5, Figure 6 and Figure 7.**

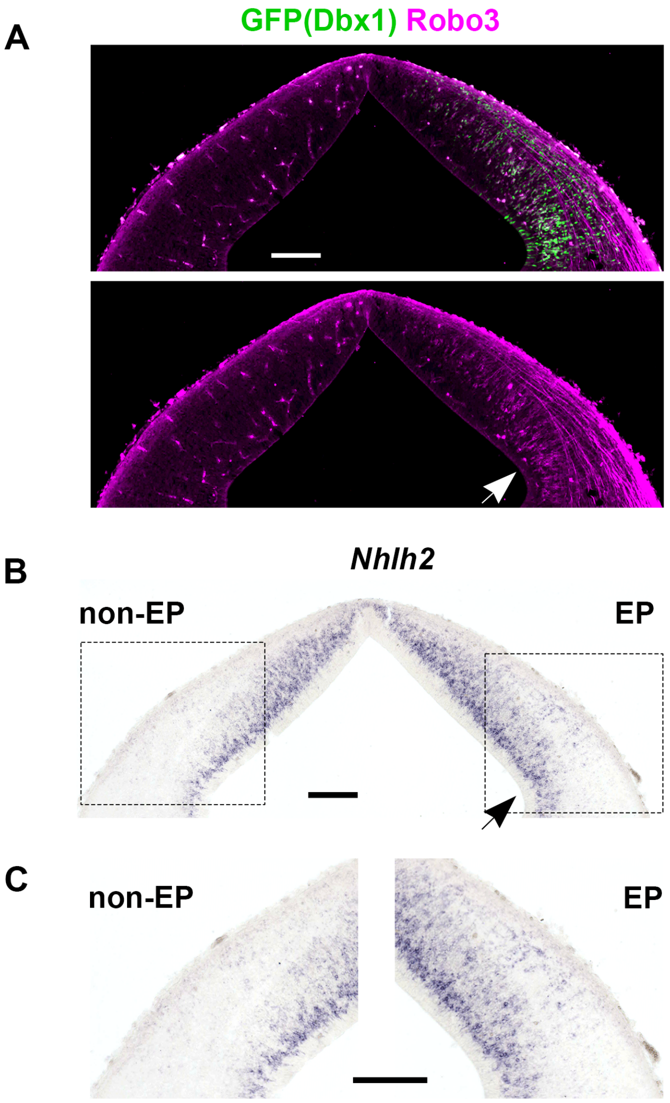


**Figure S7. Dbx1 forced expression in the midbrain at E11.5 induced ectopic expression of Nhlh2 concomitantly with Robo3.** An expression plasmid co-expressing Dbx1 and nls-EGFP was electroporated into E11.5 midbrains and samples were analyzed at E14.5 (n = 2). **(A)** Ectopic expression of Robo3 (arrow in the bottom panel) was induced by Dbx1 (GFP positive area). **(B)** ISH with a Nhlh2 probe on a section adjacent to the secion in (A). Nhlh2 IHS signal is clearly increased at the EP site (arrow) compared with the equivalent region at the non-EP side. **(C)** Higher magnificaiton images of the boxed regions in (B). We also performed ISH with the Nhlh1 probe, however, Nhlh1 ISH at E14.5 appeared diffused, hampering clear comparision between the EP and non-EP sides. Scale bars: 200 μmin (A); 200 μmin (B), 200 μm in (C). **Related to Figure 4**.

### Supplemental Table1

| REAGENT or RESOURCE | SOURCE | IDENTIFIER |
|---|---|---|
| Oligonucleotides | | |
| bHLH domain of Nhlh1 and Nhlh2 forward primer: 5'ATCTCGAGGCCACGGCCAAGT3' | This paper | N/A |
| bHLH domain of Nhlh1 reverse primer: 5'ATAGCGGCCGCTCAGACGTCCAGCA3' | This paper | N/A |
| bHLH domain of Nhlh2 reverse primer: 5'ATAGCGGCCGCCTACACGTCCAGGA3' | This paper | N/A |
| R6-bHLH domain of Nhlh1 forward primer: 5'CGCTCGAGCACTTGAGTCGTGAG3' | This paper | N/A |
| R6-bHLH domain of Nhlh2 forward primer: 5'ATCTCGAGCAGCTGAGCCGCGAA3' | This paper | N/A |
| Robo3 riboprobe forward primer: 5'ACAGCAGCCTATCTAGGCCA3' | This paper | N/A |
| Robo3 riboprobe reverse primer: 5'TCTGGTATTCAGTGATGACCCC3' | This paper | N/A |
| Nhlh1 riboprobe forward primer: 5'TGTTCAGCCACAAGCTGC3' | This paper | N/A |
| Nhlh1 riboprobe reverse primer: 5'GCGCTCCTCACGACTCAA3' | This paper | N/A |
| Nhlh2 riboprobe forward primer: 5'CTGCCAAAGGCGACTCAT3' | This paper | N/A |
| Nhlh2 riboprobe reverse primer: 5'AGACGGGGTGTTTTTGGA3' | This paper | N/A |
| Netrin-1 riboprobe forward primer: 5'CTTCCTCACCGACCTCAATAAC3' | This paper | N/A |
| Netrin-1 riboprobe reverse primer: 5'TAGAGCTCCATGTTGAATCTGC3' | This paper | N/A |
| Nhlh1-sgRNA1 forward: 5'CACCGGACCGGGCCCCGATGGTGC3' | This paper | N/A |
| Nhlh1-sgRNA1 reverse: 5'AAACGCACCATCGGGGCCCGGTCC3' | This paper | N/A |
| Nhlh1-sgRNA2 forward: 5'CACCGCTAGGTTGAAGGCTTCCACG3' | This paper | N/A |
| Nhlh1-sgRNA2 reverse: 5'AAACCGTGGAAGCCTTCAACCTAGC3' | This paper | N/A |
| Nhlh2-sgRNA1 forward: 5'CACCGCGAAGAAAAGCGCCGCCGC3' | This paper | N/A |
| Nhlh2-sgRNA1 reverse: 5'AAACGCGGCGGCGCTTTTCTTCGC3' | This paper | N/A |
| Nhlh2-sgRNA2 forward: 5'CACCGTTCTTGTCCGGAGGCAGCGT3' | This paper | N/A |
| Nhlh2-sgRNA2 reverse: 5'AAACACGCTGCCTCCGGACAAGAAC3' | This paper | N/A |
| NHLH1 mutant mice genotyping forward primer: 5'AGTCGGGCTTTAGCGACTGT3' | This paper | N/A |
| NHLH1 mutant mice genotyping reverse primer: 5'GCGAAGGCTAGGTTGAAGG3' | This paper | N/A |
| NHLH2 mutant mice genotyping forward primer: 5'CCACTACTCCTTGCAAATCAAGA3' | This paper | N/A |
| NHLH2 mutant mice genotyping reverse primer: 5'GACTAGAAAGAACAGGCTGCAAA3' | This paper | N/A |

| | | |
|---|---|---|
| NHLH2 mutant mice genotyping forward2 primer: 5’AAACTACTACCCACGCTGCC3’ | This paper | N/A |
| NHLH2 mutant mice genotyping reverse3 primer: 5’AGACGTTCTTGTCCGGAGTAG3’ | This paper | N/A |
